# The Pervasive Negative Regulation of Ion Channel Functional Families Across Human Cancers

**DOI:** 10.1101/2025.07.28.667171

**Authors:** Luca Visentin, Luca Munaron, Paola Cassoni, Luca Bertero, Alessia Andrea Ricci, Giorgia Chinigò, Federico Alessandro Ruffinatti

**Affiliations:** Department of Life Sciences and Systems Biology, University of Turin, Turin, Italy; Department of Medical Sciences, University of Turin, Turin, Italy

## Abstract

The transmembrane transport of molecules and ions is fundamental to cellular homeostasis and coordination of physiological processes. During tumorigenesis, these processes undergo significant alterations in response to oncogenic transformations and microenvironmental pressures. However, a comprehensive systems-level characterization of transportome alterations across cancer types has been lacking. Here, we integrate structural, functional, and mechanistic annotations of all known human Ion Channels and Transporters (ICTs) into a curated database, organizing them into biologically coherent gene sets based on shared physiological and biophysical properties such as permeant species, gating mechanism, and transport directionality. By leveraging Gene Set Enrichment Analysis (GSEA) across transcriptomic profiles from 19 tumor types, we reveal a recurrent downregulation of ICTs—particularly ion channels—accompanied by selective upregulation of specific pump classes. Our findings uncover a conserved signature of transportome reprogramming in cancer and provide a quantitative framework for future integrative studies of ICT function. This work highlights both the complexity and plasticity of cellular transport systems in oncogenesis and offers a resource for modeling their roles in cancer systems biology.

## 1 Introduction

In every kingdom of life, the movement of substances between the intracellular and extracellular compartments is essential for cell survival and its interaction with the surrounding environment and other cells. Apart from the free diffusion of small lipophilic molecules, these exchanges are largely mediated by transmembrane proteins of various nature.

Transport proteins can be broadly classified into two main categories (Figure 1):

- **Pores**: water-filled channel proteins that allow the facilitated passage of ions and molecules through the membrane. These can be additionally subdivided into *ion channels* proper and *aquaporins*, which mostly allow the passage of water.
- **Transporters**: membrane protein complexes that allow the passage of molecules upon conformational changes. These can be further classified into those that require ATP hydrolysis and those that do not, known as *ATP-driven* (or *primary active*) *transporters* and *SoLute Carriers (SLCs)*, respectively. A common distinction within the ATP-driven transporters is given by *ABC transporters* and *pumps*, where ABC transporters feature the conserved ATP-Binding Cassette (ABC) domain, while pumps do not.

**Figure 1:**
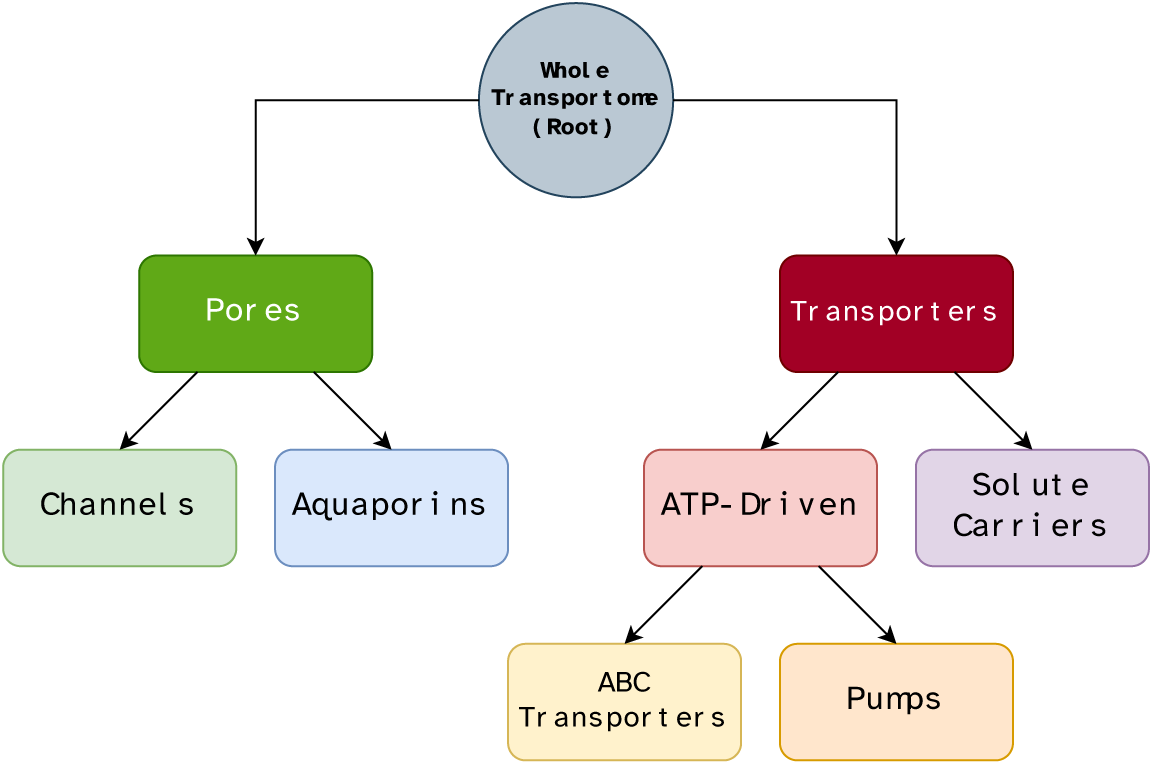
Tree of the principal classes of Ion Channels and Transporters (ICTs). Pores are water-filled channels that allow the fast movement of ions through the membrane. Aquaporins are pores specifically permeable to water. Transporters require conformational changes to exert their function, either triggered by the hydrolysis of ATP (ATP-Driven transporters) or not (Solute Carriers). The hydrolysis of ATP can be performed by the conserved ATP-Binding Cassette (ABC) domain (ABC Transporters) or not (Pumps).

Overall, the acronym “ICTs” is now commonly used to refer to the set of all the gene products that fall into one of these macro-categories, while the *-omic* layer embracing all of them is referred to as *the transportome* [1]. Currently, in humans, more than 1,000 distinct protein-coding genes are classified as ICTs or are closely associated with them.

The coordinated activity of all the components of the transportome underpins a wide array of essential physiological functions, including the maintenance of membrane potential, nutrient uptake, waste product removal, cellular signaling, and pH homeostasis across compartments. Traditionally recognized as a critical interface between cells and their environment, the transportome also constitutes one of the most important classes of drug targets in clinical pharmacology [2–4]. With the advent of high-throughput technologies and systems biology, the transportome is increasingly being studied from an omic perspective and in various physiological and pathological contexts. Not least cancer [1,5–11].

Cancer cells exhibit fundamental genetic, epigenetic, metabolic, and physiological differences from their healthy counterparts [12–14]. Manifestly, they especially differ in their relationship with the extracellular environment and other cells [15], showing a surprising degree of functional plasticity. It is fair to hypothesize that these changes may involve modifications at the level of the transportome, acting as an “adapter”— literally in the sense of “mediator of the adaptation”—of cancer cells with the tumor micro-environment [1], and at the same time ensuring the exchange of nutrients and metabolites capable to sustain the altered metabolism.

Transportome dysregulation in cancer depends, at least in part, on altered gene expression levels of the transportome genes themselves. However, as we are primarily interested in the *function* associated with ICTs, it is more informative to investigate how *classes of ICTs* with similar or identical functions change their expression levels in cancer.

A commonly employed approach to this task involves measuring gene expression in both healthy and diseased tissue samples and identifying differences through a Differential Expression Analysis (DEA). This is usually followed by some form of over-representation analysis of the resulting list of Differentially Expressed Genes (DEGs) using ontologies—such as the Gene Ontology (GO)—and statistical tests like the hypergeometric test to detect dysregulated functional categories. In studies such as ours, the significantly enriched terms must then be screened *a posteriori* to identify transportome-related terms. The effectiveness of this process depends heavily on both the statistical power of the DEA and the quality and structure of the ontology database.

An alternative, yet conceptually “reversed”, strategy is to define a limited number of gene sets *a priori*—each representing a group of functionally similar ICTs—and test them for enrichment against the differential expression data using Gene Set Enrichment Analysis (GSEA). This second approach has two main advantages. First, the weighted GSEA method accounts for the magnitude of differential expression across the gene set (i.e., the effect size), rather than simply the presence or absence of individual genes in the DEGs list. Second, the gene sets tested can be arbitrarily defined and are not constrained to a single ontology: they may be manually curated, purpose-built for a specific function or pathway, or generated by other means. Notably, given a set of gene features, one can systematically generate all conceivably meaningful gene sets and test each for enrichment.

The present work aims to profile the expression levels of transportome in the context of human cancer, not only at the level of individual ICT genes, but by extending the analysis to the higher-order level of Transporter Functional Families (TFFs). TFFs are defined as subsets of ICTs that share one or more functionally relevant features, such as ion permeability, transported solute, gating mechanism, and others. Examples of TFFs include “all sodium transporters”, “all voltage-operated channels”, “all ligand-activated and cation-permeable channels”, “all SLCs transporting organic compounds”, but also “all pumps”, or even “the whole transportome”.

We collected comprehensive information on all known genes that make up the transportome (beginning with their complete list), organized it into a structured database, and used it to systematically generate a large set of meaningful TFFs. Gene expression data for multiple tumor types and their healthy counterpart were obtained from the The Cancer Genome Atlas (TCGA) and Genotype-Tissue Expression (GTEx) databases, respectively. Protein-coding genes were ranked based on their differential expression between tumor and healthy tissues, and a pre-ranked GSEA was performed on these ordered lists to compute enrichment scores for each TFF, thereby assessing the dysregulation of functional aspects of the transportome across 19 cancer types.

Our analysis ultimately revealed that the majority of ion channel functional families—but notably not transporter families—are consistently downregulated across a broad range of malignancies, thus providing the first robust evidence of a pervasive, cancer-wide regulatory pattern specifically targeting ion channels.

To implement the analyses presented in this study, a suite of original software tools has been specifically developed. Central among these is Daedalus, a open-source and documented Python package designed to retrieve transportome-related data from various online sources and compile it in a local database *on the fly*, the Membrane Transport Protein DataBase (MTP-DB), accessible at TCP-Lab/MTP-DB. In parallel, we also provide pre-compiled database files as periodic releases. Most importantly, the entire computational workflow has been designed in accordance with the latest standards of the Open Science paradigm, prioritizing transparency, accessibility, and reproducibility. Leveraging *Kerblam!* [16], a project management and workflow orchestration technology, users are empowered to locally rerun the complete analytical pipeline—collectively named “transportome profiler” (TCP-Lab/transportome_profiler)—ensuring transparent and reproducible results.

## 2 Results

### 2.1 The Membrane Transport Protein Database

Although the GO knowledgebase [17,18] represents a valuable source of meaningful and readily available gene sets, we chose not to rely exclusively on its manually curated and automatically generated annotations. Instead, to explore all possible TFFs derived from the many transporter-related features of interest in a more unbiased and systematic way, we collected information on human transportome genes from various online databases. Then, to enhance usability and portability, we compiled all the retrieved data into a structured SQLite database, which we named the MTP-DB.

Specifically, the MTP-DB gathers information from a variety of online databases, including the Human Gene Nomenclature Committee (HGNC) database [19], the International Union of basic and clinical PHARmacology (IUPHAR) “Guide to Pharmacology” database [20], Ensembl [21], the Transporter Classification DataBase (TCDB) [22], the SLC Tables website [23], and the GO [17,18].

Particular attention in the search for information was paid to the most distinctive functional characteristics of each class of ICTs, e.g.: the solute transported and the direction of transport for SLCs, selectivity, conductance, and the gating mechanism for ion channels, as well as the pore-forming nature of the genes encoding their different subunits, and so on (see Section 5.1).

It is important to emphasize that the MTP-DB is dynamic by design, as it is automatically generated by *Daedalus*, a program capable of querying all the databases of interest and aggregating the information retrieved to rebuild the MTP-DB *on the fly* at every execution. We opted for this generation method instead of a manually updated, static database file for a few reasons: a dynamically regenerated database will always be up-to-date in respect of the upstream database changes, it does not require particular hosting capabilities to be accessed even in the far future (as the users can simply download and run the source code to obtain a copy), and it provides a clear, transparent, and traceable way to see how the source data is obtained, parsed, and stored (by examining the source code).

It is interesting to notice how information from the above upstream databases is—at the time of writing—flawed in several minor ways. For instance, the GABA_A_ receptors in the brain are ionotropic receptors notoriously permeable to chloride ions, but the HGNC database only classifies them as *ligand gated ion channels* and not as *chloride channels*. Additionally, the IUPHAR does not contain any permeability information for any of the *GABR* genes (conductance and permeability being properties of the receptor *protein complex* rather than the single subunit gene), which makes it impossible for Daedalus to classify them as permeable to chloride ions. It is also known that GABA_A_ receptors exhibit a certain permeability to HCO_3_^−^, generally in between 0.2 and 0.4 that of the chloride ions [24], but this feature is not reflected in the above databases. A similar argument applies to nicotinic acetylcholine receptors (nAChRs) and related *CHRN* subunits.

Moreover, when classifying channels by *gating mechanism*, the HGNC makes no distinction between *gating* proper (in the senses of activating stimulus) and *modulation*. As a result, some voltage-modulated channels, such as the TRP superfamily of cation channels are pooled with pure voltage-activated channels such as K_v_, Na_v_, and Ca_v_, which may not be desirable from a physiologist’s point of view. A final example is how some ion channels (including many beta subunits of potassium voltage-gated and calcium-activated channels, as well as the mitochondrial calcium uniporter family of proteins) were missing from the ion channel lists provided by the IUPHAR and the HGNC.

Even neglecting all of these considerations, the MTP-DB was still partially incomplete. For instance, when we manually inspected the list of calcium-permeable ion channels (excluding connexins), we found that (at least) 15 channels were not present in the MTP-DB, being annotated there as “cation” permeable, but without the more specific term for calcium. This means that, curiously, some well-known and biologically relevant calcium-selective ion channels (including *PIEZO1*, *HTR3A*, *P2RX7*, and some others) are not correctly annotated in any of the sources consulted by Daedalus.

All of these known defects were manually addressed in the MTP-DB through post-build hooks automatically applied by Daedalus during each build process.

In this work, the MTP-DB has been a fundamental step in collecting and organizing all the elements required for the subsequent systematic generation of the TFFs. Beyond this work, however, we hope that the MTP-DB will be an effective way to stimulate further research on Transportome by providing a centralized and human- and machine-friendly place to obtain a lot of relevant information about ICTs.

The pre-compiled MTP-DB and the Daedalus package are open-source and available for free on GitHub at github.com/TCP-Lab/MTP-DB, and we encourage readers to inspect it and propose improvements.

### 2.2 Tissue expression patterns

As a first level of investigation, we decided to collect some information on the actual presence of the different components of the transportome across the different tissues of interest.

To do this, we binarized Transcript Per Million (TPM) counts from UCSC Xena (TCGA-GTEx study) and considered only genes with a positive average log_2_(TPM + 0.001) as expressed. The gene sets of all possible TFFs were systematically generated from the MTP-DB and then filtered to mitigate redundancy according to the procedure described in Section 5.2. For each TFF and for each tissue type (cancer and healthy samples together), we computed the proportion of expressed genes out of the total number of genes in that particular TFF. The resulting expression rates are reported as a heatmap in Figure 2 and show that, while an average of ∼ 60% of all ICTs are expressed in all tissues (first row, `all_transportome`), there is a stark contrast between pores, of which fewer genes are generally expressed (∼ 30%), and transporters, which are more broadly expressed (∼ 80%).

**Figure 2:**
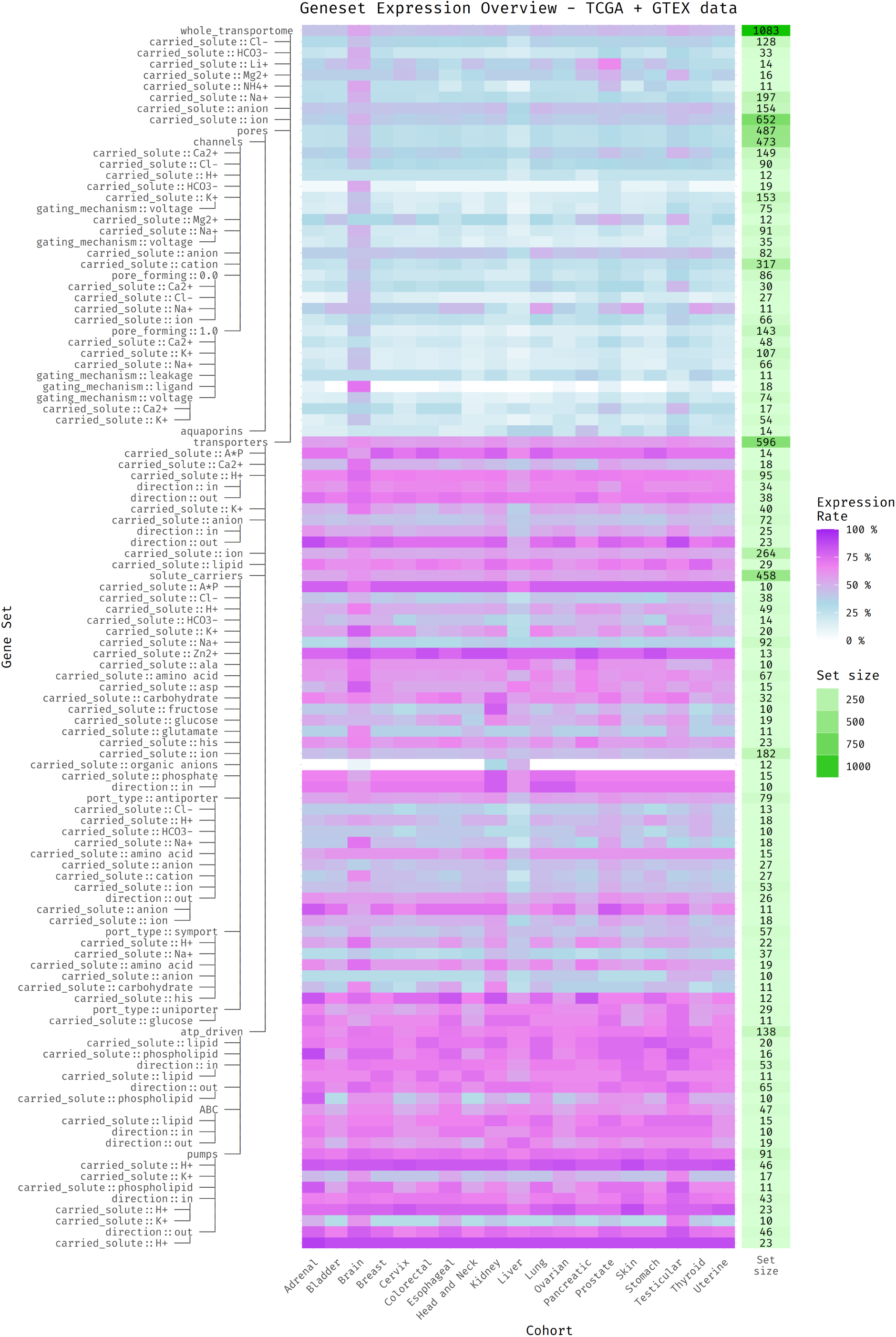
Expression rates of TFFs across tissues. Heatmap showing the proportion of expressed genes in each TFF, based on combined tumor and healthy samples from TCGA and GTEx. Gene set tree is displayed on the left, the overall size of each TFF is reported on the right (green-shaded cells), while the names of analyzed cohorts (i.e., tissue types) are reported at the bottom. While ∼ 60% of all ICTs are expressed per tissue, ion channels (pores) display lower and more variable expression (∼ 30%) compared to broadly expressed transporters (∼ 80%). Brain and testis show elevated expression across most TFFs, while liver shows a reduced diversity.

Notably, although rarely addressed explicitly—and, to our knowledge, never rigorously quantified—this observation aligns well with ICT expression patterns progressively emerging from recent omics studies [25–27]. While ion channels generally show tissue-specific and restricted expression, with only a subset active in any given tissue, SLCs, ABC transporters, and pumps show ubiquitous and broader expression across tissues.

However, strictly speaking, the fact that only a limited array of ion channels is expressed in each type of tissue does not necessarily imply that this expression pattern is highly tissue-specific (and therefore functionally characteristic). In principle, that subset of channels might really be the same (or very similar) for all tissues. This would indicate that there are some ion channels constitutively produced and shared by every tissue, together with a series of exceedingly rare channels. So, we examined the individual ICT genes expressed in the main TFFs and generated UpSet plots (Figure 3) to quantify the intersections among the different tissues in order to confirm which explanation best fit our data.

**Figure 3:**
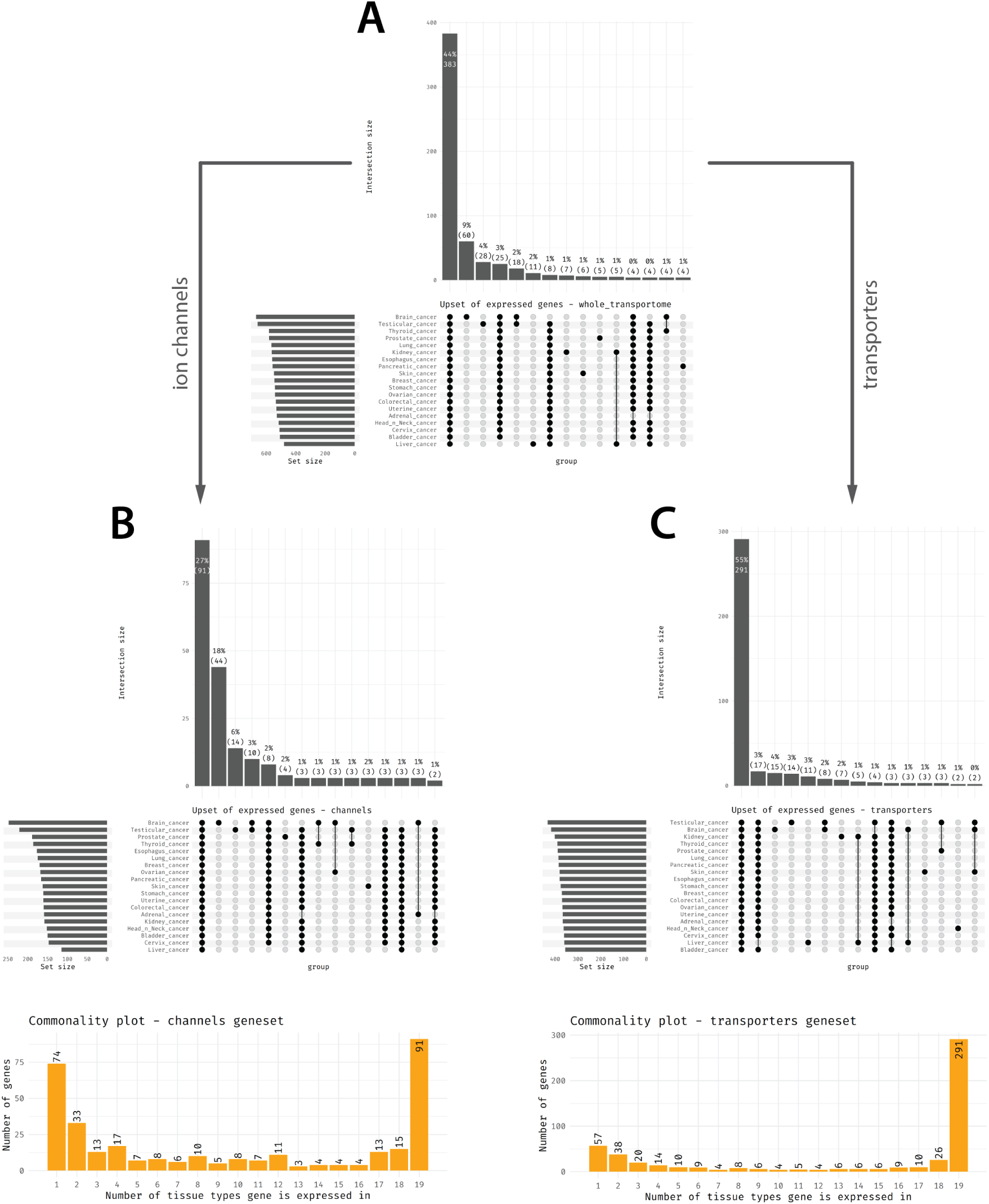
Tissue overlap of expressed genes in key Transporter Functional Families (TFFs). UpSet plots show the intersections of genes expressed across 19 tissue types for the whole transportome (A), ion channels (B), and transporters (C), based on binarized TPM expression data, considering both tumor and healthy tissues. Intersections are sorted by size, and the overlap degree is calculated as the intersection size divided by the union of all genes expressed across all tissue types (i.e., the observed spectrum for that particular TFF). Bottom panels show, respectively, how many ion channels and transporters are expressed in exactly *n* tissue types (1 ≤ *n* ≤ 19). Overall, ion channels show high tissue specificity, with only 27% expressed in all tissues and many uniquely expressed in brain or testis. In contrast, transporters are more ubiquitously expressed (55% shared), highlighting their broader physiological roles.

When considering the `whole_transportome` gene set (Figure 3, panel A), 383 genes (44% of the global observed spectrum) are shared between every tissue type, with rapidly declining intersection sizes. Tissues with the largest number of uniquely expressed ICT genes are brain (60 unique genes), testis (28) and liver (11), with all the others showing less than 10 unique genes. When considering transporters and ion channels separately, the picture changes: overall, many transporters are commonly expressed in all tissues (291, 55%, Figure 3, panel C), while the same is not true for ion channels (91, 27%, Figure 3, panel B). A large portion of channels are exclusively expressed in brain (44, 18%) or testis (14, 6%), while such uniqueness is much less pronounced when looking at transporters (15, 4% and 14, 3% respectively).

To further quantify the tissue specificity of ICT expression, we also computed the number of genes shared by an arbitrary number of tissue types (Figure 3, bottom). The resulting distribution is remarkably sharp and clearly bimodal for both ion channels and transporters, with peaks at the two extremes of the domain (i.e., genes expressed in only one tissue or in all 19). This indicates that most ICT genes are either tissue-specific or ubiquitously expressed. However, the contrast between ion channels and transporters is striking: ion channels exhibit a far more pronounced tissue specificity, with a *bin1-to-bin19* ratio of 74/91 (0.81), compared to the same ratio for transporters which is 57/291 (0.19), indicating a vast majority of genes broadly expressed across tissues.

Overall, these findings provide further quantitative support for the observation that ion channels are subject to much stricter tissue-specific regulation compared to transporters.

Looking back at the heatmap (Figure 2), we notice that, among all the tissues analyzed, brain and testes appear to be somewhat of an exception by showing a larger number of expressed genes overall (also confirmed by the UpSet plots, where tissues are ranked by the number of genes they express). Notably, this finding is also supported by the recent transcriptomic and proteomic literature [27–29]. Another irregularity, but in the opposite direction, is the liver, which appears to express a comparatively lower diversity of ion channels and transporters (being ranked last in the UpSet plots) even though, because of its unique metabolic role, it shows exceptional transporter specificity. Also this observation is compatible with current transcriptomic and functional studies [27].

Even when we look at Figure 2 by rows, we see that some ICT classes emerge as anomalies compared to the background, showing a higher degree of expression across many tissue types. In particular: anion channels, efflux anion transporters, solute carriers for Zinc (Zn^2+^) and A*P (ATP, ADP, AMP), symport solute carriers for Histidine as well as, broadly, all pumps with the exception of those for potassium (K^+^).

Remarkably, the results presented here refer to the analysis conducted by combining healthy and tumor samples within each tissue of interest. However, the same analysis conducted separately on the two cohorts yielded qualitatively similar results.

### 2.3 Shared dysregulation

As a first step in investigating cancer transportome dysregulation, we searched for individual ICTs that were most frequently altered across multiple cancer types. For all genes that we detected as expressed in Section 2.2, log_2_ FCs of Median of Ratios (MOR) normalized counts were calculated for each tissue, after pairing each TCGA tumor type to its GTEx healthy counterpart. A threshold of | log_2_ FC| ≥ 1 was then applied to generate an explorative ranking of the most shared dysregulated ICTs, the top 100 of which are shown in Figure 4.

**Figure 4:**
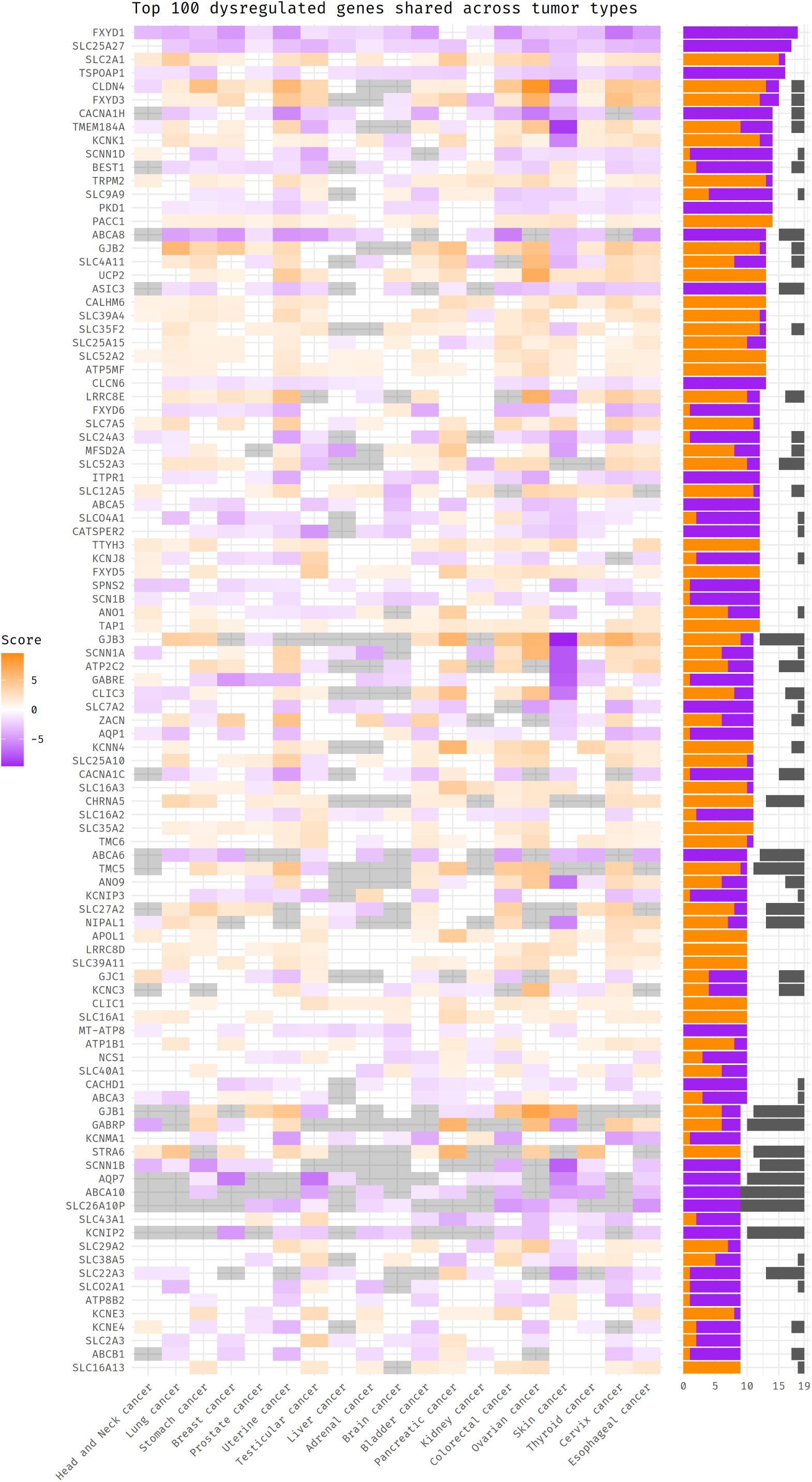
Heatmap showing genes dysregulated in multiple tumor types of the TCGA/GTEx cohort. Shown here are the top 100 genes sorted by the total number of tumor types in which they are found to be dysregulated. This number is shown on the bar plot on the side of the heatmap. Ties are broken by highest absolute average score. Purple colors represent downregulation, while orange represents upregulation. No color signifies that that gene is not dysregulated in that tumor type. A gray color signifies that that gene is not expressed in the specific tissue type, based on the results from Figure 3. This effectively reduce the maximum *potential* number of tumor types in which the gene can be dysregulated; this is reflected in the bar plot on the side. A gene is marked as either up- or down-regulated if its MOR-normalized count log_2_ FC is above 1 or below −1, respectively.

From a qualitative point of view, while an overall trend towards upregulation can be observed, it is also important to note that most ICTs show a high degree of intrinsic consistency, maintaining the same direction of alteration in all (or most) of the tumor types in which they are found to be dysregulated. This is reasonable and, to some extent, was expected as an indirect indicator of the reliability of the dataset, as it is reflective of the specific function that these genes play across different tissues and types of cancer.

Among the most commonly altered genes, *FXYD1* stands out as the single most consistently downregulated ICTs across virtually all cancer types (18 out of 19). *FXYD1* encodes phospholemman, a small membrane protein known to modulate Na^+^/K^+^-ATPase activity and regulate ion homeostasis. While its function in cancer remains understudied, recent studies have begun to implicate *FXYD1* in cancer biology, supporting its potential role in modulating proliferation and cell signaling [30,31]. Other members of the FXYD family, including *FXYD3*, *FXYD5*, and *FXYD6*, are also represented in the heatmap, although with lower ranks and different dysregulation patterns, confirming their diverse and context-dependent roles in cancer (see e.g., [32–34]).

The second most dysregulated ICT belongs to the SLC25 family of mitochondrial solute carriers. Overall, four SLC25 genes—*SLC25A27* (UCP4), *SLC25A8* (UCP2), *SLC25A15* (ORNT1), *SLC25A10* (DIC)—are present in our top-100 gene list. *SLC25A8* and *SLC25A27* (also known as mitochondrial uncoupling proteins 2 and 4, respectively) separate oxidative phosphorylation from ATP synthesis and are implicated in mitochondrial membrane potential regulation and protection against oxidative stress. *SLC25A15* encodes the mitochondrial ornithine transporter 1, essential for the urea cycle and nitrogen metabolism, while *SLC25A10* dicarboxylate carrier mediates the exchange of dicarboxylates and inorganic phosphate, supporting biosynthetic pathways and redox balance. Each of these has already been documented, to varying degrees, as playing a role in cancer metabolism and progression of some tumor types [35,36]. Our analysis, however, extends the domain of validity—and therefore the relevance—of these observations.

Beyond mitochondrial solute carriers, many other SLCs appear in the heatmap of Figure 4, in line with existing literature linking dysregulation of many solute carriers to all hallmarks of cancer [37]. In particular, the most prominent cluster is that linked to glucose and energy production, which includes *SLC2A1* (GLUT1), *SLC2A3* (GLUT3), and monocarboxylate transporters *SLC16A1* (MCT1), *SLC16A2* (MCT8), *SLC16A3* (MCT4), and *SLC16A13* (MCT13). These transporters are central to the metabolic reprogramming of cancer cells, supporting the enhanced uptake of glucose and the efficient export of glycolytic byproducts such as lactate, distinguishing features of the Warburg effect. Consistently, *GLUT1* and *GLUT3* are frequently overexpressed in tumors to sustain high glycolytic flux [38], while *MCT1* and *MCT4* facilitate lactate shuttling, contributing to the acidic microenvironment that favors invasion and immune evasion [39]. Furthermore, *SLC4A11*—another highly ranked carrier in our gene list—may also contribute to the altered pH dynamics observed in cancer. *SLC4A11* functions as an acid extruder and its upregulation in tumors is likely to represent an adaptive response to aerobic glycolysis to sustain intracellular alkalinization while promoting extracellular acidification [40], a combination known to support tumor progression, invasion, and metastasis [41–45].

As for the ion channels most consistently dysregulated across cancer types, we identified members from a wide range of families. The voltage-gated calcium channels *CACNA1H* (T-type Ca_V_3.2 subunit) and *CACNA1C* (L-type Ca_V_1.2 subunit) have already been reported to be dysregulated in numerous types of cancer, leading to aberrant calcium signaling [46,47]. The potassium leak channel *KCNK1* (K_2P_1.1, TWIK-1) contributes to membrane potential regulation and its over-expression has been implicated in tumor cell survival [48]. The epithelial sodium channel *δ* subunit (*δ*ENaC, as encoded by *SCNN1D*) is still poorly studied and characterized [49], but, interestingly, is one of the most commonly downregulated ion channel gene in our dataset. On the contrary the *α* ENaC (encoded by *SCNN1A* and also present in our top-100 list) is known to be involved in sodium absorption and to play a role in some cancer types, such as breast [50].

Another top-ranked gene in our analysis is the cation channel *TRPM2*, which mediates calcium influx in response to ADP-ribose (ADPR) production following oxidative stress and has been associated with cell survival, DNA damage response, and resistance to therapy in multiple cancer types [51]. It is interesting to note that, despite the abundance of literature on the role of TRP channels in cancer, *TRPM2* is the only TRP channel in the ranking of the 100 most commonly dysregulated ICTs (although if we look at the list of the 100 most dysregulated *ion channels*, rather than the entire transportome, we also find *TRPV2*, *TRPV4*, *TRPV6*, *TRPC1*, *MCOLN2*, *MCOLN3*; see supplementary figure v). This could indicate a greater specificity of TRPs compared to other classes of ion channels, which directly reflects in their alteration profile.

Among ion channels, a special role is played by connexins, which form gap junctions between cells. In our analysis we found that *GJB2* and *GJB3* (also known as *connexin 26* and *connexin 31*, respectively) are upregulated in most cancer types. *GJB2* is reported to be involved in cancer stemness, drug resistance, and metastasis in many solid tumors [52,53], including breast [54,55], pancreas [56], kidney [57], and lung [58]. Likewise, although *GJB3* is not among the most studied connexins, recent works seem to confirm its pan-cancer dysregulation [59] and its dysfunctional role in lipid metabolism by supporting cancer cell energy needs and membrane synthesis [60]. In addition, *GJB1* (*connexin 32*) and *GJC1* (*connexin 45*), two other connexins among the best known in the literature for their role in cancer, also appear in our top-100 list.

The most commonly altered ABC family member is *ABCA8*, which, in our dataset, is consistently downregulated in almost all cancer types in which it is expressed. *ABCA8* is involved in lipid metabolism regulation and, unlike other ABCs that are often associated with drug resistance, it is increasingly recognized as a putative tumor suppressor in multiple cancer types [61,62]. Similar findings have been reported for other ABCs belonging to the same sequence homology cluster—*ABCA5*, *ABCA6*, *ABCA10*—which also appear in the highest positions of our shared dysregulation heatmap. Thus, our analysis confirms these observations and broadens them to a wider scope.

More generally, Figure 4 recapitulates many well-established findings on ion channels and transporters in cancer, thereby validating both the reliability of the original dataset and the robustness of our analytical pipeline. At the same time, it extends these observations to a broader pan-cancer context and uncovers novel dysregulated ICTs whose involvement in tumorigenesis has not yet been reported, offering new avenues for future investigation.

### 2.4 Gene Set Enrichment Analysis

To extend the resolution of our analysis beyond the single-gene level, we aimed to advance our research to a more integrative and conceptually meaningful layer, able to capture coordinated patterns of regulation among functionally related ICTs. For this purpose, we employed the MTP-DB and the same TFF gene sets already introduced in Section 2.2 to perform a systematic GSEA. This allowed us to quantify directional and statistically significant shifts in the expression of entire ICT families across tumor types, thereby capturing higher-order regulatory trends potentially masked at the single-gene resolution.

For each gene and each TCGA/GTEx tissue pair, the Cohen’s *d* of the log_2_ MOR-normalized counts was calculated and used as a metric for the pre-ranked GSEA. The resulting Normalized Enrichment Scores (NESs) scores are plotted in Figure 5. Similar figures generated with all the other considered metrics are available in Supplementary Section 1.

**Figure 5:**
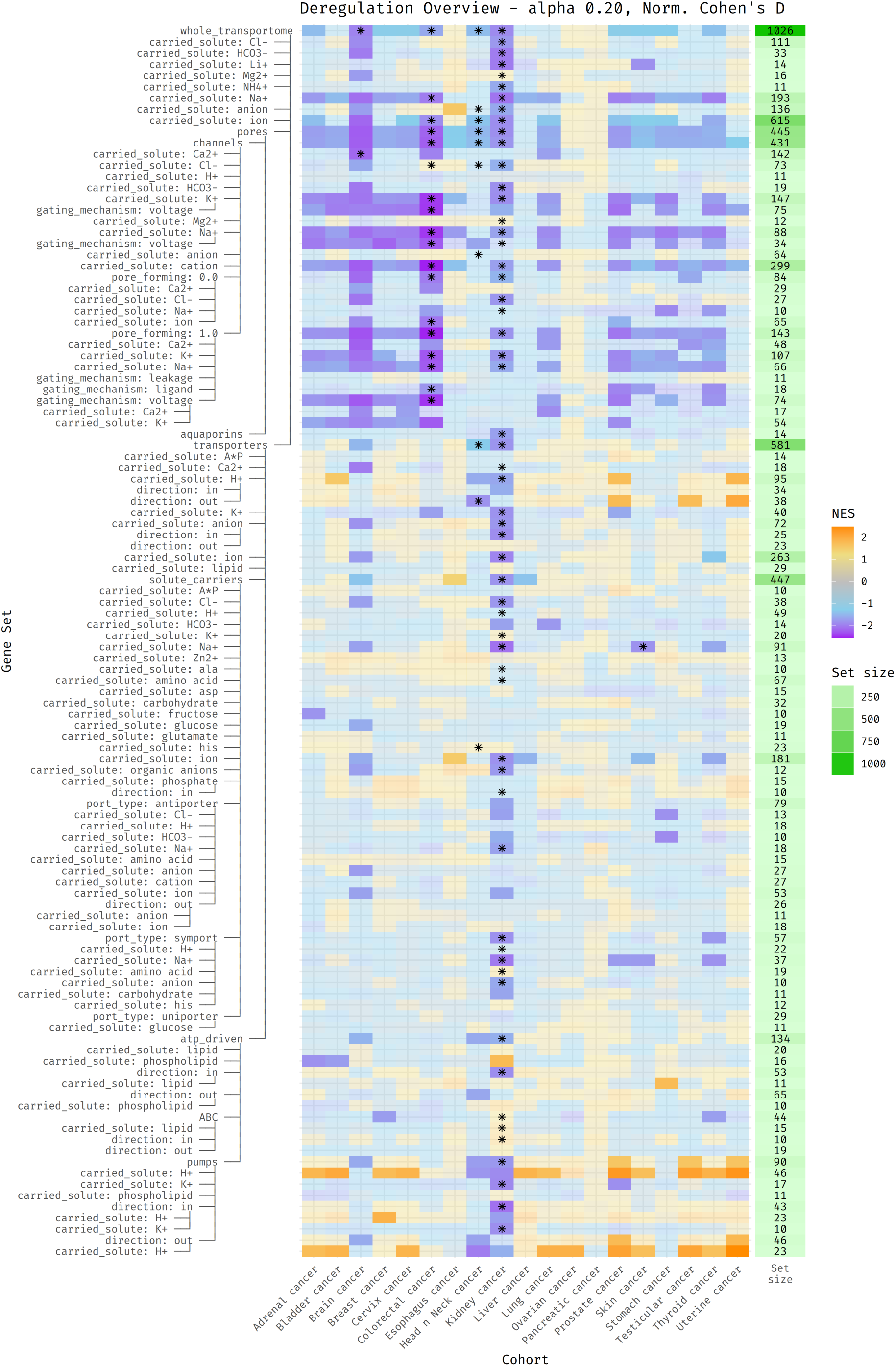
Heatmap of GSEA NESs for all TFFs derived from the MTP-DB, given TCGA and GTEx differential gene expression. Gene lists were sorted based on Cohen’s *d* scores between tumor samples and controls. These scores were computed over expression matrices normalized with the Median of Ratios (MOR) method as implemented by `pydeseq2`. Blue/purple colors denote downregulated gene sets, while yellow/orange denote upregulation. A faded color represents non-significant *p*-values with reference to a significance threshold of *α* = 0.20. Asterisks at the center of each cell represent significant results if GSEA is performed with absolute fold change values instead of the regular signed one, as a metric of “general dysregulation” of that gene set. Columns are sorted alphabetically.

As shown in the heatmap, when considering the transportome as a whole (first row), 11 out of 19 cancer types exhibit a significant global alteration in their ICT configuration and, crucially, this dysregulation is consistently negative. This trend becomes even more striking when ion channels and transporters are analyzed separately. In particular, a pervasive downregulation of ion channels is observed in 16 of the 19 cancer types, indicating a widespread suppression of the biological processes they mediate. In stark contrast, only 3 cancer types show a statistically significant dysregulation of the `transporters` gene set, which is also downward in all cases.

Overall, we believe the most salient and consequential finding of our study is this “double asymmetry”: at the gene set level, ion channels—but not transporters—are broadly altered across most cancer types, and this alteration is consistently in the direction of downregulation, suggesting a cancer-wide attenuation of some functions typically mediated by ion fluxes, such as signal transduction, volume regulation, and electrochemical communication.

Looking more closely at the `transporters` gene set, a noteworthy exception to the general trend emerges: several ATP-driven transporters, particularly H^+^-extruding pumps, show consistent upregulation across multiple cancer types. This observation is strongly supported by existing literature and aligns well with the established view of cancer cell adaptation to metabolic reprogramming [39] as well as what already said in Section 2.3 about the increased expression of *SLC4A11* and proton-coupled monocarboxylate transporters [37,40]. Tumor cells often rely on aerobic glycolysis, leading to the accumulation of acidic byproducts such as lactate. In this context, the upregulation of proton pumps can be interpreted as an adaptive mechanism to maintain intracellular alkalinity while acidifying the extracellular milieu. Such pH modulation not only supports continued glycolytic flux and cellular proliferation but also contributes to tumor progression by facilitating immune evasion, invasion, and metastasis [41–44].

Interestingly, within the ion channel gene sets (upper part of the heatmap), the down-regulation appears to predominantly affect the pore-forming subunits (`pore_forming: 1.0`), while auxiliary/regulatory components (`pore_forming: 0.0`) are largely spared, thus suggesting a targeted suppression of the core functional machinery responsible for ion conduction. Notably, the most strongly downregulated TFFs include voltage-gated sodium and potassium channels, which play essential roles in controlling membrane excitability, cell volume regulation, and signal transduction, with well documented implications in cancer [63].

Looking at the global dysregulation of the different cancer types, pancreatic, ovarian, and liver cancers are the least dysregulated categories, while kidney, brain, and colorectal cancers seem to be the most dysregulated tumor types, with a pronounced downregulation for kidney cancer. Notably, while most tumors show widespread alterations primarily in ion channels, brain and (even more so) kidney cancer also display a significant and pervasive dysregulation of transporter categories, including SLC. This broader dysregulation may reflect a deeper metabolic and homeostatic rewiring, in line with the unique physiological demands of these tissues and the special set of ICTs they express. For instance, renal carcinomas are known to undergo profound metabolic reprogramming and disruption of epithelial transport processes [64], while glioblastomas show extensive transcriptomic remodeling of ion transport and metabolism-related genes, often associated with tumor invasiveness and resistance to therapy [65].

To validate these findings from the TCGA/GTEx cohort, we also analyzed 15 independent single-study datasets from Gene Expression Omnibus (GEO), which benefit from more specific tumor–normal pairings and are free from possible cross-cohort batch effects, though their smaller sample sizes result in markedly reduced statistical power. The results of the same analytical pipeline, previously applied to the TCGA-GTEx data and now conducted on the 15 GEO datasets, are shown in Figure 6.

**Figure 6:**
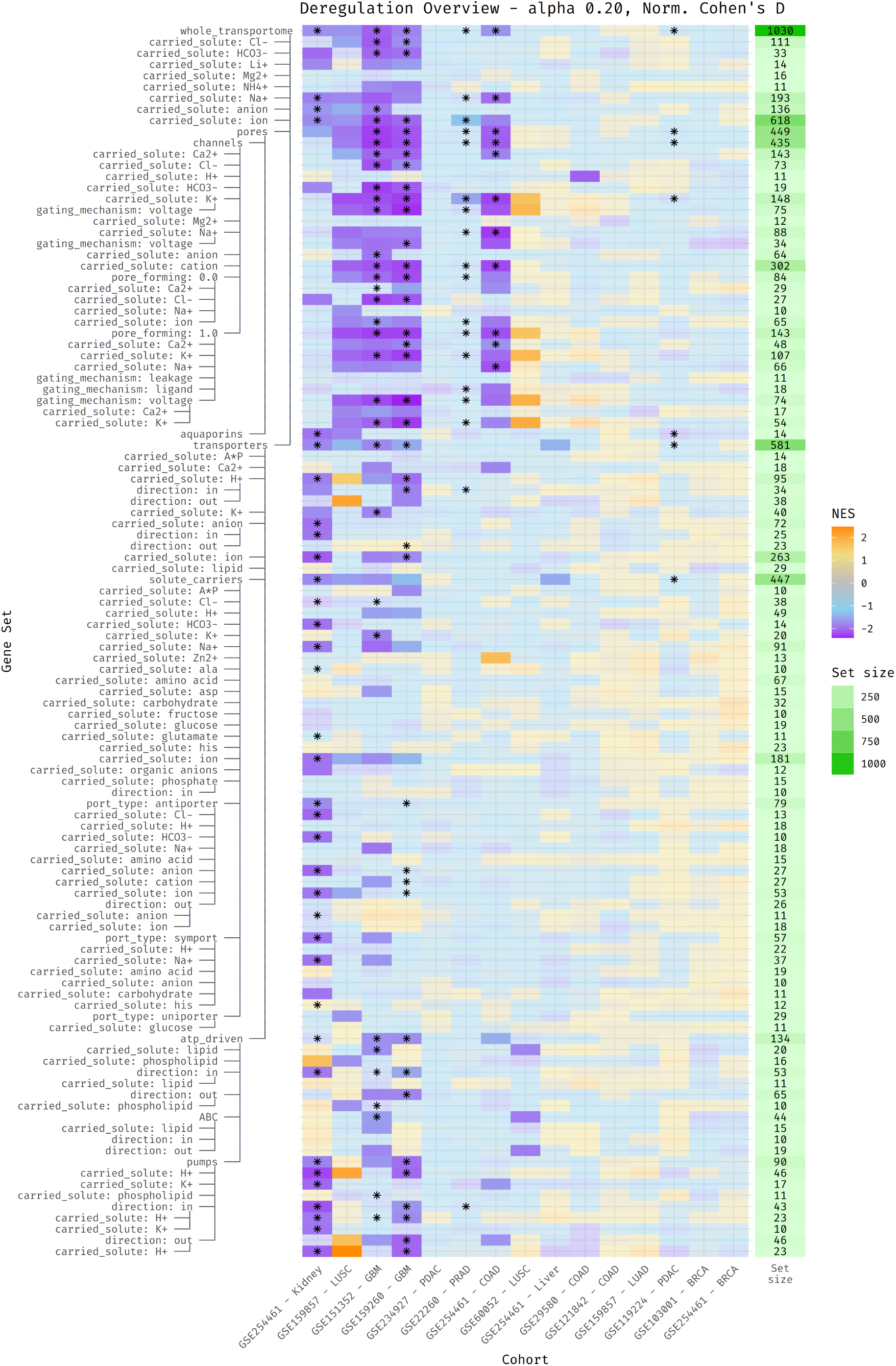
Heatmap of GSEA NESs for all TFFs derived from the MTP-DB, given the differential gene expression analysis of the GEO cohorts. Data were analyzed using the same procedure employed for the TCGA-GTEx cohorts, the results of which are shown in Figure 5. Briefly, expression matrices were first normalized using the MOR method, then Cohen’s *d* scores calculated between tumor samples and controls were used to sort gene lists. Blue/purple colors denote downregulated gene sets, while yellow/orange denote upregulation. A faded color represents nonsignificant *p*-values (i.e., above the significance threshold *α* = 0.20). Asterisks at the center of each cell represent significant results if GSEA is performed with absolute fold change values.

Despite some exceptions, this second analysis broadly confirms the main trends observed in the TCGA-GTEx cohort. Both glioblastoma (GBM) studies—GSE151352 and GSE159260—closely replicate the signature of the widespread downregulation of ion channel functional families observed in the larger dataset. Similarly, one of the three colon adenocarcinoma studies (GSE254461 - COAD) shows a consistent dysregulation pattern, while the other two (GSE29580 and GSE121842), which include only 6 and 8 samples respectively, fail to detect significant alterations, possibly due to limited statistical power. Among the two lung squamous cell carcinoma studies, one (GSE159857 - LUSC) reproduces the TCGA-GTEx signature with remarkable fidelity, including the upregulation of proton pumps, while the other (GSE60052) shows only sparse and inconsistent dysregulation. Conversely, the lung adenocarcinoma dataset analyzed (GSE159857 - LUAD) did not display any significant enrichment scores. Kidney cancer in the GEO cohort (GSE254461 - Kidney) mirrors the TCGA-GTEx profile overall, though with a curious attenuation of the strong downregulation of ion channel TFFs seen in the larger dataset. Prostate adenocarcinoma (GSE22260 - PRAD) confirmed the dysregulation profile only when the absolute-value metric was applied. Pancreatic cancer (GSE234927 and GSE119224 - PDAC) appears weakly dysregulated in both cohorts, while liver (GSE254461 - Liver) and both breast cancer studies (GSE103001 and GSE254461 - BRCA) did not show any significant shifts in transporter functional families.

Importantly, even with these discrepancies, most GEO studies tend to confirm the over-arching trend of ion channel downregulation. In cases where results diverge, possible variability in sample or sequencing quality and/or the small sample sizes combined with general cancer heterogeneity of the GEO datasets are the most likely explanation.

## 3 Discussion

Our analysis reveals a pervasive and cancer-wide dysregulation of the transportome, and of ion channels in particular, with a striking functional asymmetry relative to other transporter families. While solute carriers and most ATP-driven transporters remain largely unaffected—or in some cases, upregulated—ion channels are consistently repressed across the majority of tumor types. Importantly, the fact that downregulation of ion channels is shared by most tumor types suggests some form of mechanistic rationale for this phenomenon.

### 3.1 Expression patterns reflect physiological functions

A key observation from our data is the contrasting expression patterns between ion channels and other transporter classes across healthy and tumor tissues as shown in Figure 2 and Figure 3. Ion channels tend to show tissue-specific and restricted expression, with only a subset active in any given tissue, whereas SLCs, ATP-driven pumps, and ABC transporters are broadly and ubiquitously expressed. This dichotomy mirrors their prevalent functional repartition: signal transduction for channels, and metabolic and homeostatic support for transporters.

Ion channels are intrinsically suited for fast and precise signal transduction (see e.g., [66]). Their capacity for rapid gating in response to voltage changes, ligands, or mechanical stimuli enables the tight spatiotemporal control of ion fluxes. A canonical example is intracellular calcium signaling: Ca^2+^ dynamics (oscillations, spikes, or sustained elevations) are essential for decoding a wide variety of extracellular and intracellular cues in both normal physiology and disease [67–69].

Ion channels also form multimeric complexes with variable subunit composition, enabling considerable functional diversity and sensitivity to modulation by auxiliary proteins, drugs, or second messengers. This structural modularity provides an evolutionary substrate for fine-tuning signal transduction in a tissue-specific manner [70]. While there are many channels for—say—calcium transport, only a small subset of them can properly respond to the stimuli that a specific cell needs to be sensible to (voltage, mechanical stretch, specific intra/extra-cellular messengers), often in combination with cell-type-specific signaling cascades.

In contrast, solute carriers and pumps are responsible for essential, constitutive functions shared by virtually all cell types. They mediate the uptake of nutrients (e.g., glucose, amino acids), removal of metabolic waste (such as monocarboxylates), regulation of intracellular pH, and maintenance of ion gradients (as in the case of the homeostatic return to initial conditions following ion channel opening) [25,26]. These transporters operate on slower timescales than channels, but with the ability to transport substances against a concentration gradient through direct or indirect consumption of metabolic energy. This made them the perfect candidates for evolution to act upon when selective pressure favored organisms that could absorb or excrete molecules for metabolic or functional reasons. It follows that transporters supporting these basic cellular functions would be widely (co-)expressed [25] and—as we shall see—more resilient to changes in expression in pathological situations such as cancer.

In synthesis, transporters’ features make them ideal when supporting cell metabolism and those of ion channels make them suitable for signal transduction, but their physiological roles are not interchangeable. As we already noted, this separation of functions is nicely reflected in the expression patterns highlighted by Figure 2.

An additional observation that support these different physiological roles of transporters and ion channels comes from the brain, where we notice a marked increase in the number of expressed ion channels, and the liver, where the opposite is true (Figure 2 and Figure 3). The brain is possibly the organ in which signal transduction is most central to its physiological role, and this is reflected in an increase in the number of expressed ion channels. A similar trend is seen in the testes and prostate, which also show elevated ion channel expression, likely related to sperm maturation, endocrine regulation, or neural innervation. In contrast, the liver, where metabolic function dominates, shows relatively low expression of ion channels but retains abundant—and tissue-specific— expression of SLCs and pumps.

### 3.2 Physiological functions are selectively affected by cancer

Framing ion channels as signal transducers and transporters as metabolism-supporters allows us to make sense of the results of the GSEA of Figure 5. The metabolic reprogramming of tumor cells may favor the retention of “housekeeping” transporter families (or the upregulation of those supporting nutrient uptake and pH control), while the tissue-specific signaling functions of ion channels may become dispensable—or even detrimental—under oncogenic selection pressures.

In other words, the fact that cancer cells behave very differently from normal cells implies changes in the way extracellular and intracellular signals are transduced. Accordingly, a general and selective dysregulation of ion channels, directly reflecting the altered behavior of cancer cells, would be expected, which is precisely what we observed across most cancer types. What is surprising, however, is that this selective dysregulation manifests specifically as a systematic *downregulation*, a pattern that emerged clearly only when analyzing functional families (i.e., TFFs) rather than individual genes.

Some reasons for this strong evidence are here hypothesized. First, cancer cells might generally be selected to become less sensitive to external stimuli. This makes sense when considering that cancer cells are resistant to many extracellular signals such as those that would trigger apoptosis, cell-to-cell contact proliferation inhibition or the switch to an inflammatory phenotype. Thus, “silencing” of cellular responsiveness may be a key element in the reprogramming that underlies tumorigenesis, representing a mechanism through which cells adopt a less interactive, more autonomous phenotype optimized for unchecked proliferation and survival in a hostile microenvironment. For instance, mechanosensitive and calcium-permeable channels, which facilitate responses to physical and metabolic stress, are repressed in multiple cancers, enabling resistance to apoptosis and promoting migration and survival under adverse conditions [34]. Additionally, membrane depolarization—often a consequence of reduced K^+^ channel activity—has been associated with proliferative and undifferentiated phenotypes, and oncogenic signaling [63,71].

More generally, we could say that signal transduction is essential in cell cooperation and the “core mandate” of system-over-cell survival that characterizes all multicellular organisms. When this “mandate” is broken, as in the case of cancer cells, suppression of signal transduction is an expected outcome.

Secondly, as signal transduction is a key feature of healthy tissue and cell identity, the marked dedifferentiation of cancer cells towards less specialized cell phenotypes—as in the case of the Epithelial-Mesenchymal Transition (EMT) observed in carcinomas—might involve the downregulation of signal-transducing, tissue-type-specific ion channels. Voltage-gated channels, for instance, are commonly expressed in excitable or differentiating cells and are often downregulated in tumors as part of a broader dedifferentiation process [63].

Such a pattern aligns with the hypothesis that cancer cells dampen their sensitivity to extrinsic stimuli as a strategy to evade differentiation cues, apoptotic triggers, or contact inhibition signals, all of which are often mediated through ion flux.

Thirdly, changes in cell shape, volume, and volume-to-surface ratios due to active proliferation might also be reflected in the reduced transcription of ion channels, as energy is diverted toward cell growth and duplication rather than toward other less “pressing” physiological functions.

Interestingly, Ko and colleagues [72], albeit through a less systematic approach, also reported a bias toward downregulation of many families of ion channels in highgrade glioma, even if they didn’t observe this trend in other cancer types [73,74]. This was probably because their approach, although “channelome-oriented”, was based on single-gene expression, rather than functionally related gene families, as in the case of the GSEAs we performed.

As a corollary to all these considerations, if our observation of widespread downregulation of ion channel TFFs across multiple tumor types reflects a broader reprogramming of signal transduction whereby cancer cells progressively dedifferentiate and lose sensitivity to external cues, a similar trend could also be observed at the *receptome* level. A recent paper [75] systematically mapped the co-regulation of G Protein-Coupled Receptors (GPCRs) with their ligands and biosynthetic enzymes across a large transcriptomic pan-cancer dataset. Their *omics* approach was very close to the one adopted in our work, at least from a conceptual point of view. In their analysis, Arora *et al* identified extensive downregulation of GPCR signaling axes in specific cancer subtypes, particularly in association with mutations in tumor suppressor genes and dedifferentiated phenotypes such as brain and prostate tumors. While their study did not explicitly frame this downregulation as a loss of sensory input, the coordinated suppression of both receptors and upstream ligand-producing enzymes strongly suggests a reduction in extracellular signal responsiveness. Thus, we could speculate that the concurrent downregulation of two major classes of transduction proteins, GPCRs and ion channels, may reflect a shared strategy by which tumors disengage from tissue-specific signaling constraints and acquire a more plastic, self-sustained, and signal-decoupled phenotype. However, in contrast to our observations, Arora and coworkers also reported many GPCR-ligand or GPCR-enzyme axes as systematically co-upregulated, depending on the tumor subtype. It would be interesting to investigate whether, also in the case of receptors, it is possible to establish a relationship between receptor (family) specificity and the direction of their dysregulation.

More generally, when considering the receptome layer, other evidence suggests a more fundamental and possibly complex rewiring of how cancer cells interpret environmental cues, which is not limited to the mere desensitization. For instance, the role of the nervous system in cancer progression has recently gained attention, particularly with regard to GPCRs and ionotropic receptors, both in CNS [76] and outside [77].

The pervasive downregulation of ion channel functional families observed in our analysis does not necessarily contradict the widely held view that ion channel activity can be upregulated to support cancer cell proliferation and migration. In fact, at the single-gene level, our data also show a bias toward upregulation for several individual channels Figure 4. However, when the analysis is elevated to the functional family level, a different picture emerges and ion channel downregulation becomes the prevailing trend, revealing a previously underappreciated dimension of cancer-associated molecular remodeling.

At the same time, our data show that not all TFFs are equally affected. A notable example is the consistent upregulation of H^+^-extruding ATPases, which supports the hypothesis of active pH regulation as a central adaptive feature of cancer metabolism. This finding is in line with extensive prior work on pH dynamics in tumors [41–45], and validates the utility of our TFF-based approach in capturing physiologically meaningful shifts in gene regulation.

Despite prior reports indicating dysregulation of several transportome components in pancreatic cancer [78,79], an intriguing finding of our study is the apparent lack of significant enrichment for ICT gene sets in PDAC from both TCGA/GTEx and GEO cohorts, thus suggesting some intrinsic biological or structural complexity for this tumor type, rather than a mere artifact of the TCGA/GTEx pipeline. While multiple technical and biological factors could underlie this observation (including sample heterogeneity, gene set composition, or metric sensitivity) we focus here on two biological explanations that are particularly relevant. First, the absence of clear enrichment may result from divergent regulation within ICT families. Pancreatic tumors are known for their metabolic plasticity [80], and it is plausible that the selective upregulation of specific transporters is counterbalanced by the repression of others, thus averaging out enrichment signals when considered at the level of entire transporter families. This hypothesis seems to be supported by our secondary analysis of the GSE119224 GEO dataset revealing a significant dysregulation of the main TFFs (`whole_transportome`, `channels`, `aquaporins`, `solute_carriers`) when absolute-value metric is used. Second, the complex tumor microenvironment of pancreatic cancer, characterized by pronounced desmoplasia and a high proportion of stromal and immune cells [81], may mask epithelial-specific transcriptional changes in bulk RNA-seq data. This effect is particularly relevant for ICTs with low expression or restricted cellular localization, whose dysregulation may remain undetected when signal is diluted by the abundant non-malignant stromal compartment.

Overall, transportome dysregulation may be the result of several different characteristics of cancer and further research is required to determine these underlying mechanisms. However, our findings suggest a selective evolutionary or mechanistic pressure against ion channel function in tumorigenesis, opening new avenues for the functional dissection of ion flux modulation in cancer biology.

### 3.3 Limitations and Future Perspectives

While our study presents a robust framework for the systematic exploration of the transportome in cancer, several limitations must be acknowledged, many of which offer fertile ground for future developments.

First, a significant portion of the project effort was devoted to the design and implementation of the Membrane Transport Protein DataBase (MTP-DB), which serves as the foundation for the generation of Transporter Functional Families (TFFs) and subsequent enrichment analyses. However, despite integrating data from multiple curated sources (e.g., IUPHAR, HGNC, GO [17–20]), known inaccuracies and misclassifications persist within upstream databases and are likely to propagate downstream to the TFFs and their enrichment scores. Although we implemented several post-build hooks to correct the most prominent inconsistencies (see Section 2.1), a full manual curation remains desirable to improve classification reliability.

Such a manual curation step could also enrich the database with additional informative features currently absent, such as subcellular localization (e.g., plasma membrane vs. intracellular membranes), ligand types (e.g., ions vs. peptides), or functional distinctions between activating and modulating stimuli. These enhancements would allow the definition of novel TFFs reflecting underexplored functional roles and improve the granularity of future analyses.

Another conceptual limitation stems from the structural complexity of many ICTs. Several ion channels and pumps operate as multimeric complexes composed of poreforming, regulatory, and auxiliary subunits. Examples include homo- and heteromeric TRP channels [82,83], GABA_A_ receptors [24], and various ATPases [84]. In these cases, essential functional properties like conductance and gating arise from the assembled protein complex rather than individual gene products. As such, functional inference at the single-gene level is inherently limited [70]. We believe that addressing this point would require the development of a new ontology accounting for ICT complexes and subunit-level dependencies. Since this is an endeavor that is far beyond the scope of the present work, here we adopted the widely popular simplification that each individual ICT is identified with the gene of one or more subunits it consists of, with the implicit assumption that component genes in a multimeric complex are expressed in concert, properly assembled, and targeted in the “right” place.

Additionally, our investigation (like others of the same kind [75]) is based on transcriptomic data, which, while widely available and analytically tractable, does not fully capture protein function. For ICTs to be operational, gene transcription must be followed by translation, proper folding, membrane targeting, and, where needed, complex assembly. Proteomic approaches would be more suited to assess functional ICT expression, yet these remain technically and logistically more challenging, with limited publicly available large-scale datasets [1]. Other aspects of cell transcriptomics could also be informative, and might potentially be integrated in our analysis to provide additional insight on the transportome remodelling in cancer. For example, reconstructing which transcription factors are linked with the dysregulation patters we detected could shed light on the phenomenon from a network level. Epigenetic modifications, such as gene and enhancer methylation states, might also be involved in these patterns, and could be a fertile area for additional investigation.

Finally, our analysis did not stratify tumors by stage, gender, or histological subtype. While these variables undoubtedly influence gene expression profiles and may reveal more specific patterns of ICT dysregulation, our decision to treat each tissue as a single, unified category was motivated by two primary considerations. First, our aim was to generate a broad, pan-cancer overview of transportome remodeling that highlights general trends and principles governing ICT dysregulation, rather than focusing on more granular context-specific effects. Second, many TCGA/GTEx tissue comparisons suffer from unbalanced sample sizes when further subdivided, reducing statistical power and robustness of downstream GSEA analyses. GEO cohorts were used as a secondary check as they represent controlled, small-scale analyses for which we can more accurately combine tumor and healthy samples, also to confirm the validity of TCGA/GTEx macro-scopic pairings. Nevertheless, future studies may explore whether the dysregulation of specific TFFs correlates with clinical or biological covariates, particularly in the case of cancers with strong hormonal components (e.g., breast, prostate, or ovarian cancer), or those with established molecular subtypes, provided that large, well-annotated cohorts are available.

## 4 Conclusion

In this work, we present the first comprehensive, pan-cancer analysis of the human transportome, revealing a consistent and selective downregulation of many ion channel functional families across the majority of tumor types, in contrast to the relative stability of solute carriers and the context-dependent upregulation of specific ATP-driven pumps, particularly proton extruders.

Emerging as a new hallmark of cancer, the systematic attenuation of the channelome —often associated with cell-type specific signaling functions—potentially reflects dedifferentiation and reduced sensitivity to external cues, supporting autonomous growth and survival. On the contrary, metabolic and homeostatic transport functions are preserved or reinforced.

These findings were enabled by a fully reproducible computational pipeline centered around the Membrane Transport Protein DataBase (MTP-DB)—a dynamically generated and manually curated resource integrating multiple annotation sources—and a recursive strategy for the programmatic generation of thousands of non-redundant Transporter Functional Families (TFFs). Future extensions should also integrate proteomic data and multimeric protein ontologies to fully capture the complexity of ICT remodeling in disease. Importantly, both the generation of gene sets and the analysis of associated expression data are entirely data-agnostic, and the same transportome profiling strategy demonstrated here for cancer can be readily applied to any omics dataset.

The transportome plays a central role in cell physiology, is a major target of available drugs, and its plasticity is essential when the extracellular environment changes. Altogether, the systematic patterns observed here point to a previously underappreciated dimension of tumor physiology, laying the groundwork for future functional investigations of ion transport in cancer biology.

## 5 Materials and Methods

### 5.1 The Membrane Transport Protein Database

The MTP-DB was designed as a universal repository for collecting information on ICTs currently available on the web from the different consortia involved in various ways in the annotation and characterization of human Transportome genes.

A Python package nicknamed *Daedalus* was created to download, parse, and compile all this information in a portable SQLite database, thus recreating the MTP-DB *on the fly* at every execution. In particular, to populated the MTP-DB, Daedalus extracted data from the HGNC database [19], the IUPHAR “Guide to Pharmacology” database [20], Ensembl [21] through the use of BioMart [85], the TCDB [22], the SLC Tables website [23], the Catalogue Of Somatic Mutations In Cancer (COSMIC) [86] database, and the GO [17,18]. At the time of writing, due to licensing issues, the COSMIC data were omitted from the latest version of the MTP-DB and ignored in this work. Parsing logic still exists for such data, however, should licensing change in the future. The specific data downloaded from each database and the methods used to retrieve them can be found in Table 1.

**Table 1:** List of accessed databases, methods of access, and the use of the downloaded data in the MTP-DB. API access or other formalized methods were preferred, if available. If not available, the raw HTML page was heuristically processed.

| Database / Repository | Access method | Retrieved Data |
| --- | --- | --- |
| HGNC Database | FTP repository, direct download URLs | Transporter group classification, Gene Names and Symbols (indirectly) |
| IUPHAR / Guide to Pharmacology | Direct download URLs | Transporter group classification, Ion Channel selectivity and conductance metrics, protein ligand information, ligand-to-protein interactions |
| Ensembl (BioMart) | BioMart XML requests | Transporter group classification, Gene Names, Gene Identifiers (Ensembl Genes, Transcripts and Protein IDs, RefSeq Gene and Protein IDs, PDB IDs) |
| TCDB | Direct download URLs | Transporter group classification |
| SLC Tables (BioParadigm) | Webpage Scraping | Solute carriers selectivity information |
| Gene Ontology | BioMart XML requests | Ion Channel selectivity information |

In addition to the information from these sources, manual interventions were made in the form of post-build hooks injected by Daedalus at the end of the build process to correct defects, errors, and omissions. Notably, even such “terminal patches” can be easily inspected in detail by the reader in order to get an idea of the degree of manual curation within the MTP-DB. If needed, post-build hooks can be turned off to obtain a version of the database free of manual annotations.

As we believe that the MTP-DB can still contain flaws and would benefit from additional manual curation, we set up the GitHub repository with a collaborative approach in mind, following Open Science practices. This would allow researchers that notice an issue with the annotations in the database to participate in the creation of a more useful resource. A complete list of the current known issues that affect the database can be found at [this link](https://github.com/TCP-Lab/MTP-DB/issues).

An overview of the database schema, with information on the type of data in many of the main tables, can be seen in Table 2.

**Table 2:** Brief description of the data contained in the main tables present in the MTP-DB.

| Table name | Description |
| --- | --- |
| gene_ids | Genome-wide gene identifiers from Ensembl (“ENSGs”) and their versions. |
| transcript_ids | Transcript identifiers from Ensembl (“ENSTs”), and their version, associated with gene identifiers. |
| mrna_refseq | RefSeq transcript identifiers linked with other identifiers. |
| protein_ids | Protein identifiers from Ensembl (“ENSPs”) associated with their respective ENSTs, PDB accession codes and RefSeq protein IDs. |
| gene_names | ENSGs associated with their respective HUGO gene IDs, symbols and names. |
| IUPHAR_targets | Every gene considered as a “target” by the IUPHAR, with relevant names, IDs and family information. |
| IUPHAR_ligands | Every ligand considered by the IUPHAR, as well as its type, name, ENSG (if relevant), pubchem SIDs and CIDs, and information on drug status. |
| IUPHAR_interaction | Every interaction between a IUPHAR-recognized ligand and human protein target, with relevant information. |
| TCDB_ids | The entirety of the TCDB linked with the respective ENSPs, split by type, subtype, family, subfamily and superfamily. |
| channels | All genes catalogued as “channels” by IUPHAR or the HGNC, with information on their gating mechanism, carried solute and relative and absolute conductance. Information from GO regarding permeability information is also included. |
| aquaporins | All aquaporin genes as well as their expression tissue. |
| solute_carriers | All genes catalogued as as “solute carriers” by IUPHAR or the HGNC, as well as their carried solute, rate of transport, stoichiometry, port type, direction, and more. |
| pumps | All active transporters (excluding ABC transporters), correlated with their carried solute, rate, transport direction, net transported charge, and more. |
| ABC_transporters | All ABC transporter genes, correlated with their carried solute, rate, direction, and more. |

To support a more varied search strategy, especially when querying data for SLCs, a sort of internal thesaurus was implemented, adding synonyms or more general terms to the database besides the already-present information on the carried molecule in all relevant tables. For example, we inserted the more general values of “cation” and “anion” next to each particular ion species; we added more general synonyms such as “carbohydrate” for mannose and glucose; “amino acid” for all basic amino acids; and so on. The thesaurus is also consulted to correct any parsing errors made by Daedalus, and manually addresses some imperfections in the source data (e.g., different codes for the same amino acid are all converted to the three-letter code). The thesaurus is contained in a single, human-readable comma-separated-values file, available in the remote repository for manual inspection. It is worth noting that the thesaurus is applied multiple times on the source data, until all the necessary conversions are done.

Table indexes are automatically created by Daedalus in order to speed up data lookup. Thanks to these indexes, even complex queries involving multiple tables are completed almost instantaneously.

Additionally, note that the data blob downloaded and then used by Daedalus to generate the database is locally saved as a Python Pickle file, and also distributed at every release, and is the only file that the package needs to regenerate the database. This can be useful to exactly reproduce the database in the future even after the remote databases have updated.

The database is versioned using the CalVer specification, with representation `MAJOR.YY.0W[_MINOR][-Modifier]`. At the time of writing, the latest version of the database is `1.25.24`, featuring a total of 1083 ICT gene entries (473 ion channels, 14 aquaporins, 47 ABC transporters, 91 pumps, and 458 solute carriers).

The pre-compiled MTP-DB release (in the form of the generated SQLite file) and the Daedalus package are open-source and available for free on GitHub at www.github.com/TCP-Lab/MTP-DB. Periodically, a published version of the database will be released on GitHub also deposited on Zenodo at the following DOI: <u>10.5281/zenodo.12622364</u>, for reproducibility purposes. The latest MTP-DB release can be downloaded directly from [this URL](https://github.com/TCP-Lab/MTP-DB/releases/latest/download/mtpdb.sqlite.gz) in GZIP-compressed SQLITE format.

### 5.2 Generation of TFF gene sets

In this work, a TFF is defined as any subset of transportome genes representing a precise group of ICTs that share one or more functionally relevant characteristics among those included in the MTP-DB.

To systematically generate all the possible TFFs, a Python script nicknamed *Ariadne* was created to recursively query the database file produced by Daedalus. Resulting TFFs were structured as a tree in which each node represents a separate TFF. All transportome genes are present in the root node, while subsets of the parent nodes are considered in every child node.

To organize the genes into meaningful TFFs, we seeded the algorithm to always produce a manually defined tree—or “core” tree—reflecting the classification schema already shown in Figure 1. This subdivision purposefully follows also the internal schema of the database, since most nodes represent a table or a combination of whole tables. The algorithm then retrieves all relevant information for each node of the core tree and finds all the possible sub-classes based on the available data, finally appending each subclass to the tree as a child node. This approach is then repeated for each added child, recursively creating smaller and smaller child gene sets, stopping when a child node cannot be meaningfully subdivided further.

This iterative process generates a very large number of gene sets (with the current version of the MTP-DB, this number is ∼ 6000) which are highly congruent with each other. For this reason, a filtering procedure on the resulting tree was implemented, pruning away leaf nodes that are highly similar with other nodes in the tree. Node similarity was measured by the Sørensen–Dice coefficient *κ* as defined below:

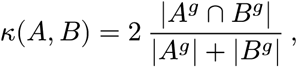

where *A* and *B* are nodes in the tree of TFFs *T*, associated to the sets of genes *A*^*g*^ and *B*^*g*^, respectively, and |*X*| represents the number of items in the set *X*. *κ* is a value ranging from 0 to 1, where 1 means perfectly congruent nodes, and 0 meaning completely different nodes. Given a tree *T*, we calculated the similarity index for each node *A* as

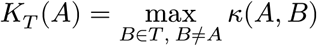

If *K*_*T*_ (*A*) is greater than a user-defined threshold value *η* (generally close to 1), the node *A* is deemed redundant and pruned.

We consider for pruning only leaf nodes. This is to prevent deletion of a branch due to the redundancy of a node close to the root and therefore potentially losing meaningful downstream child nodes. After all leaves of the tree are considered for pruning, if the algorithm pruned at least one leaf, it is executed again to further reduce the size of the tree.

This process returns a tree in which all leaf nodes are sufficiently different from each other. On the contrary, internal nodes are always preserved, even in the presence of high similarity scores between them, since they cannot be eliminated without the removal of meaningful leaf nodes.

For efficiency purposes, the algorithm does not consider data columns that contain too many missing values (deeming them not meaningful to split on, with the default being > 50% emptiness) and does not generate gene sets that are smaller than a threshold (by default 10 genes).

The specifications for the core tree, the SQL calls used to retrieve the data for each node in the core tree, the threshold for *κ*, the pruning direction, and other settings are all easily configurable as command-line arguments. This makes this approach easily adaptable to future versions of the MTP-DB and even other kinds of SQLite and more generally SQL-based databases provided by other researchers. For our run, we used the default parameter values, while *κ* was set to 0.88.

### 5.3 Input expression data

Most of the analysis pipelines presented here have been applied to the expression data made available by the UCSC Xena project [87] through the so-called Toil recompute [88] in which ∼ 20, 000 RNA-seq samples from different consortia—among which TCGA and GTEx—were reanalyzed to create a consistent meta-analysis free of computational batch effects.

In particular, the TCGA/TARGET/GTEx cohort was selected and, after excluding TARGET samples, RSEM expected raw counts were used as direct input for DEA and as a starting point to compute gene pre-ranks for GSEA. Expression data in TPMs were also downloaded from the same resource and used for expression rate computation.

The entire TCGA/TARGET/GTEx study is available both directly from the UCSC Xena platform ([here](https://xenabrowser.net/datapages/?cohort=TCGA%20TARGET%20GTEx&removeHub=https%3A%2F%2Fxena.treehouse.gi.ucsc.edu%3A443)) and from a re-upload we made (with permission from the original authors) on Zenodo for long-term preservation (https://zenodo.org/records/10944168). In the analysis workflow presented here, the Zenodo clone is used.

To supplement the TCGA/GTEx analysis, further independent studies from the GEO archive were selected for a number of tumor types. The list of such studies as well as their accession numbers can be found in Table 3. If a single study encompassed multiple tumor types, it was split up following such types and the samples of interest were processed independently.

**Table 3:**
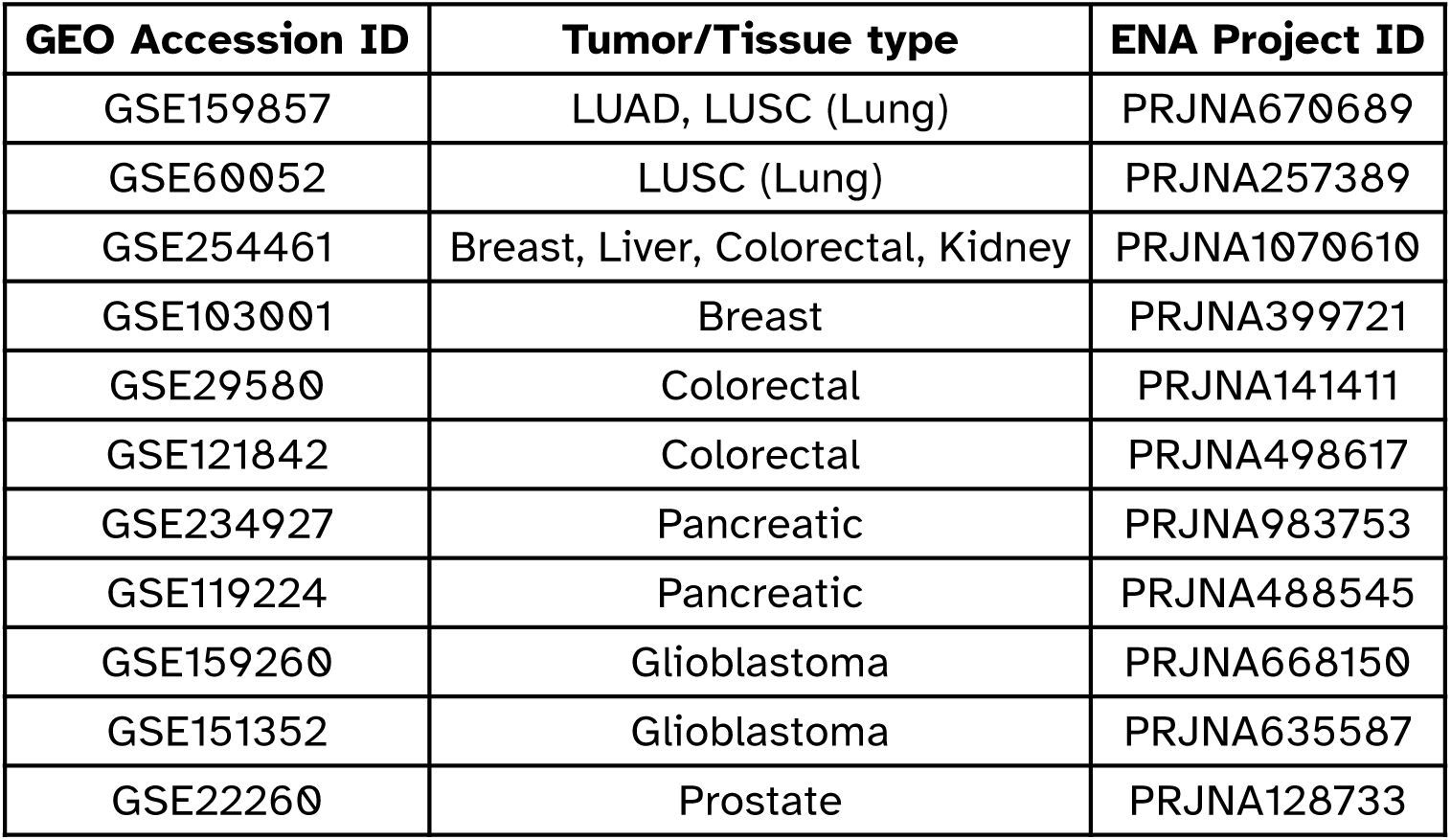
GEO accession IDs of selected GEO studies used in the analysis. Highlighted are the tumor or tissue types of the studies as well as the correlated European Nucleotide Archive (ENA) project IDs used to derive metadata for each study.

| **GEO Accession ID** | **Tumor/Tissue type** | **ENA Project ID** |
| --- | --- | --- |
| GSE159857 | LUAD, LUSC (Lung) | PRJNA670689 |
| GSE60052 | LUSC (Lung) | PRJNA257389 |
| GSE254461 | Breast, Liver, Colorectal, Kidney | PRJNA1070610 |
| GSE103001 | Breast | PRJNA399721 |
| GSE29580 | Colorectal | PRJNA141411 |
| GSE121842 | Colorectal | PRJNA498617 |
| GSE234927 | Pancreatic | PRJNA983753 |
| GSE119224 | Pancreatic | PRJNA488545 |
| GSE159260 | Glioblastoma | PRJNA668150 |
| GSE151352 | Glioblastoma | PRJNA635587 |
| GSE22260 | Prostate | PRJNA128733 |

Data were harvested directly from European Nucleotide Archive (ENA) and analyzed with a standard pipeline involving trimming with BBDuk, alignment with STAR [89], and quantification with RSEM [90] as implemented in the `x.FASTQ` tool [91]. Options were kept at their defaults for all the analysis to homogenize this processing step. The same tool was used to harvest metadata for the samples in a standardized format.

The final output of the realignment work, as well as the harvested metadata for GEO samples, is available on Zenodo at the following DOI: <u>10.5281/zenodo.11183835</u>. Data used in the pipeline are directly downloaded from the above archive. More information on this realignment pipeline is also available at the above page.

RSEM “expected count” expression data was normalized following the Median of Ratios (MOR) normalization procedure by using the same function implemented in the DESeq2 package [92,93].

If *c*_ij_ is the read count for gene *i* in sample *j* (out of *m* total samples)

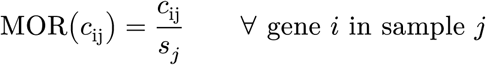

where *s*_*j*_ is the *size factor* for sample *j*, calculated as

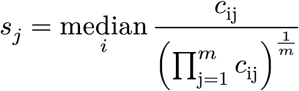

In log scale this is equivalent to mean-centering rows/genes and then median-centering columns/samples, which makes the bulk of non-DEGs similar across each sample.

### 5.4 Determination of expression rates

To determine if a given gene was expressed or not in a set of samples, TPM counts from UCSC Xena were used, as detailed above. A gene was deemed to be expressed in a set of samples if its mean log_2_(TPM + 0.001) across all samples, including both healthy and tumor samples, was greater than 0. In linear terms, a gene was considered to be expressed if its average TPM count was greater than 0.999 [94,95].

The rate of expression of the genes in a TFF gene set within a specific group of samples was defined as the number of genes that are actually expressed in that set of samples over the size of the entire gene set.

To evaluate the degree of overlap among all tissue and tumor types in terms of expressed genes in a given TFF, UpSet plots were created for three salient gene sets—`whole_transportome`, `channels`, and `transporters`—and the degree of overlap was calculated for each intersection as the size of the intersection divided by the size of the largest theoretical intersection, that is, the size of the union of all genes expressed across all tissue types. These plots were generated using the UpSetR R package [96].

### 5.5 Pairing of healthy and diseased tissues

To perform DEA and GSEA on the TCGA/GTEx cohort, TCGA tumor expression data and their GTEx “healthy” counterparts needed to be paired for each organ/tissue type separately.

Sample pairing was based on the macroscopic tissue- or organ-level grouping provided by UCSC Xena as a single integrated metadata file.

To subset the very large expression matrix file based on Xena metadata, a Python package called *metasplit* was implemented, which makes use of the low level Rustcompiled xsv tool to speed up the computation [97].

GEO cohorts were used as a secondary check of the results obtained from the TCGA/ GTEx analysis, as they represent controlled, small-scale analyses for which tumor and healthy samples can be more accurately combined.

The numerosity of each comparison is available in Table 4 for both TCGA and GEO samples.

**Table 4:** Sample sizes of both TCGA/GTEx and GEO datasets, separated by tumor type or GEO accession and by status, e.g. Tumor samples or Healthy controls.

| Tissue | Tumor | Control |
| --- | --- | --- |
| Adrenal | 78 | 127 |
| Bladder | 408 | 29 |
| Brain | 690 | 1154 |
| Breast | 1100 | 292 |
| Cervix | 306 | 99 |
| Colorectal | 290 | 349 |
| Esophagus | 183 | 666 |
| Head And Neck | 521 | 45 |
| Kidney | 887 | 158 |
| Liver | 372 | 161 |
| Lung | 1014 | 398 |
| Ovarian | 428 | 89 |
| Pancreas | 180 | 172 |
| Prostate | 496 | 152 |
| Skin | 470 | 557 |
| Stomach | 414 | 211 |
| Testicular | 155 | 166 |
| Thyroid | 513 | 339 |
| Uterine | 239 | 102 |
| GSE103001 | 23 | 23 |
| GSE119224 | 7 | 7 |
| GSE121842 | 4 | 4 |
| GSE151352 | 13 | 13 |
| GSE159260 | 16 | 6 |
| GSE159857 (LUAD) | 16 | 15 |
| GSE159857 (LUSC) | 18 | 13 |
| GSE22260 | 21 | 11 |
| GSE234927 | 4 | 4 |
| GSE254461 (BRCA) | 22 | 3 |
| GSE254461 (COAD) | 6 | 6 |
| GSE254461 (KI) | 6 | 4 |
| GSE254461 (LIV) | 4 | 4 |
| GSE29580 | 3 | 3 |
| GSE60052 | 80 | 8 |

### 5.6 Computation of dysregulation metrics

To observe the dysregulation of single genes, a DEA was performed on the RSEM expected counts for every tissue type. The downstream GSEA analysis differs significantly based on which dysregulation metric is selected as a basis for it (see Section 5.7), therefore we needed to compute several different dysregulation metrics.

We implemented a Python package nicknamed *gene_ranker* which can compute all the dysregulation metrics that we inspect. In particular, we obtained the following statistics: Log Fold Change, Cohen’s *d*, DESeq2 shrunken log fold change (through the usage of `pydeseq2` [92,93]), Baumgartner-Weiss-Schindler test statistic [98] (as implemented in `scipy`) and the signal to noise ratio. All input data were normalized just before computing the measures with the MOR procedure as previously detailed (see Section 5.3).

Given the very high numerosity of the considered samples (Table 4), we opted to not calculate *p*-values associated with any metric, and instead rely on the calculated statistics directly. We assume that any change, however small, would generate a significant *p*-value, and therefore it is more meaningful to filter the data based on the threshold on the metric’s value instead.

When referring to single genes, as in Figure 4, we deemed as “upregulated” or “down-regulated” those that had a log_2_ Fold Change above 1 or below −1, respectively.

### 5.7 Geneset enrichment analysis

Zyla, J., Marczyk, M., Weiner, J. & Polanska, J. [99] showed that the usage different GSEA metrics has dramatic effects on the outcome of the computation. For this reason, we tested all metrics computed by `gene_ranker`, as detailed above, in order to compare them and select the most meaningful. Confirming the results cited above, the choice of metric to used had dramatic effects on the computed GSEA Normalized Enrichment Scores (NESs).

GSEA was performed in the “pre-ranked” mode with input data the ranked gene lists of a given metric, and the set of all genesets generated from the MTP-DB. To run GSEA, we used an R implementation of the algorithm by Korotkevich, G. *et al.* [100] in the package `fgsea`. The family-wise error rate was corrected at the level of every tumor/ healthy comparison type, in other words, at every tumor type.

Overall, we believe the most representative and easy-to-interpret result is the one obtained by running GSEA with the Cohen’s *d* metric. Other metrics either provide very few interpretable signals (e.g. BWS Test Statistics) or are difficult to interpret after GSEA (e.g. DESeq2-Shrunk Fold changes). In any case, all calculations were performed with all metrics. The full set of results may be appreciated in Supplementary Section 1.

## 6 Resource availability

### 6.1 Lead contact

Requests for further information and resources should be directed to and will be fulfilled by the lead contact, Federico Alessandro Ruffinatti.

### 6.2 Materials availability

This study did not generate new unique reagents.

### 6.3 Data and code availability

We leveraged Kerblam! as project management tool [16]. This allows the reader to directly obtain all data and code from the Kerblam! replay package that is available from <u>10.5281/zenodo.16355762</u>. The previous link also contains instructions on how to obtain all results presented in the paper leveraging Kerblam!. A containerized execution environment is available on Docker Hub at [this URL](https://hub.docker.com/layers/cmalabscience/transportome_profiler_env/3/images/sha256-3e80d20d91ec4f8718514864485fa2d6813943c119ed5666118338417432eed0) and is fetched automatically by Kerblam! upon replay.

#### Data

This paper analyzes existing, publicly available data. TCGA/GTEX data recomputed by XENA is accessible at [this URL](https://xenabrowser.net/datapages/?cohort=TCGA%20TARGET%20GTEx&removeHub=https%3A%2F%2Fxena.treehouse.gi.ucsc.edu%3A443) or, equivalently, from Zenodo at zenodo.org/records/10944168. Raw GEO data can be accessed from the Gene Expression Omnibus (available at https://www.ncbi.nlm.nih.gov/geo/) with the GEO IDs visible in Table 3, and the results of our realignment and quantification, used in the presented analysis, can be found at the following DOI: <u>10.5281/zenodo.11183835</u>. The MTP-DB release used in this work can be found on Zenodo at <u>10.5281/zenodo.15632999</u>. All figures, relevant tables, and other results have been deposited in Zenodo with DOI <u>10.5281/zenodo.15281114</u>.

#### Code

All original code has been deposited on Zenodo with the following DOI <u>10.5281/zenodo.15281353</u>, and is available on GitHub at the following URL <u>TCP-Lab/transportome_profiler</u>. To produce the results presented in this work, we used the repository at version 2.2.

#### Additional information

Any additional information required to reanalyze the data reported in this paper is available from the lead contact upon request.

## 7 Acknowledgements

The results published here are in part based upon data generated by the TCGA Research Network: https://www.cancer.gov/tcga. We thank the TCGARN for their precious contribution.

## 8 Author Contributions

L.V.: Conceptualization, Formal analysis, Methodology, Software, Visualization, Writing original draft; L.M.: Supervision, Funding acquisition, Writing - review & editing; P.C.: Conceptualization, Writing - review & editing; L.B. Conceptualization, Validation, Writing review & editing; A.A.R: Writing - review & editing; G.C.: Writing - review & editing; F.A.R.: Conceptualization, Formal Analysis, Investigation, Methodology, Software, Supervision, Validation, Writing - original draft.

## S.1 Alternative GSEA metrics

These plots in this section show all metrics tested during the analysis in the same heatmap styles shown in the main text, with the exception of those already in the main text.

While the different patterns of dysregulation of channels, transporters and pumps are apparent when running GSEA with all metrics, the fine values change dramatically.

**Supplementary Figure G:**
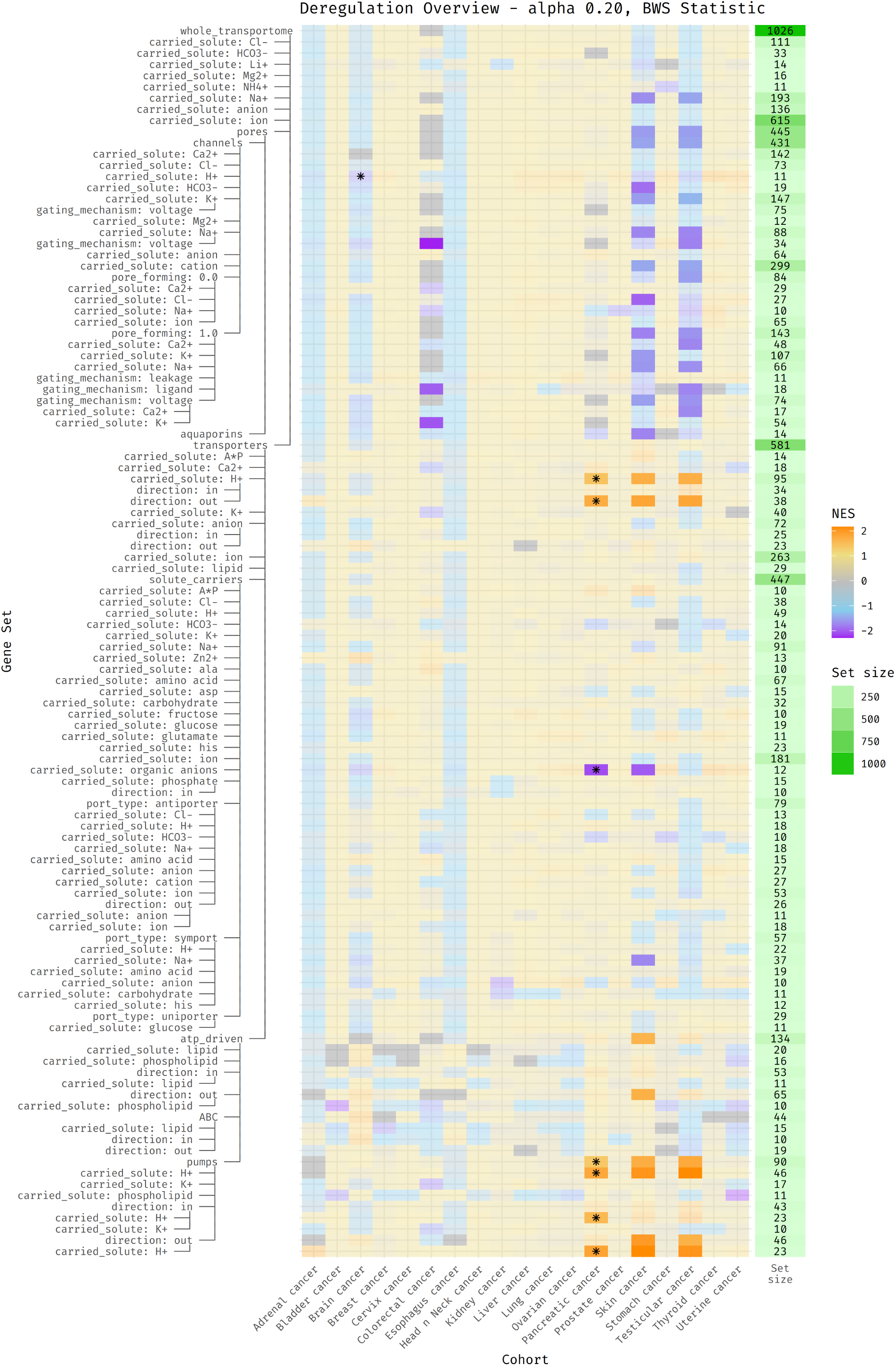
Supplementary deregulation heatmap with metric “bws_test”.

**Supplementary Figure H:**
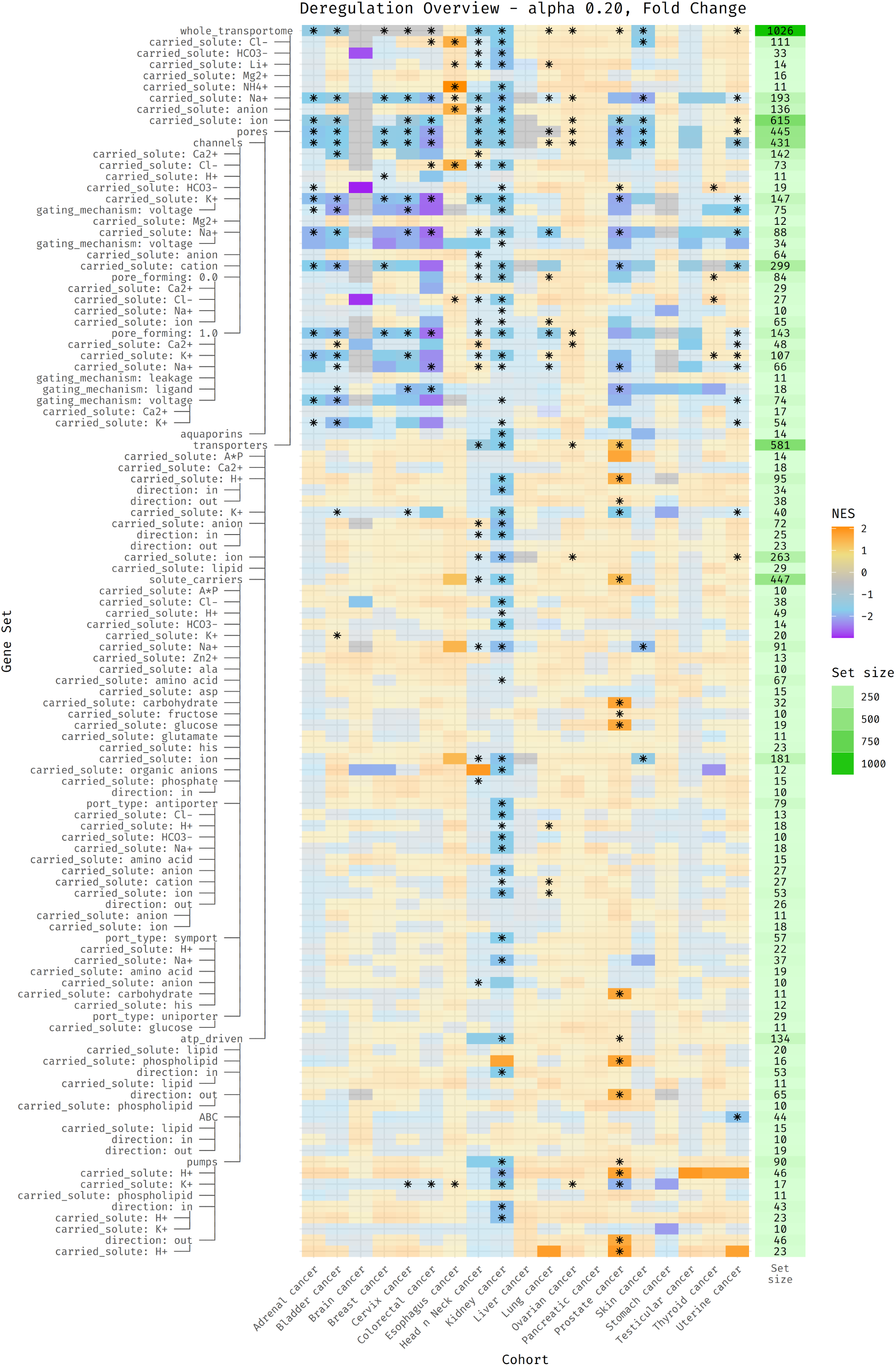
Supplementary deregulation heatmap with metric “fold_change”.

**Supplementary Figure I:**
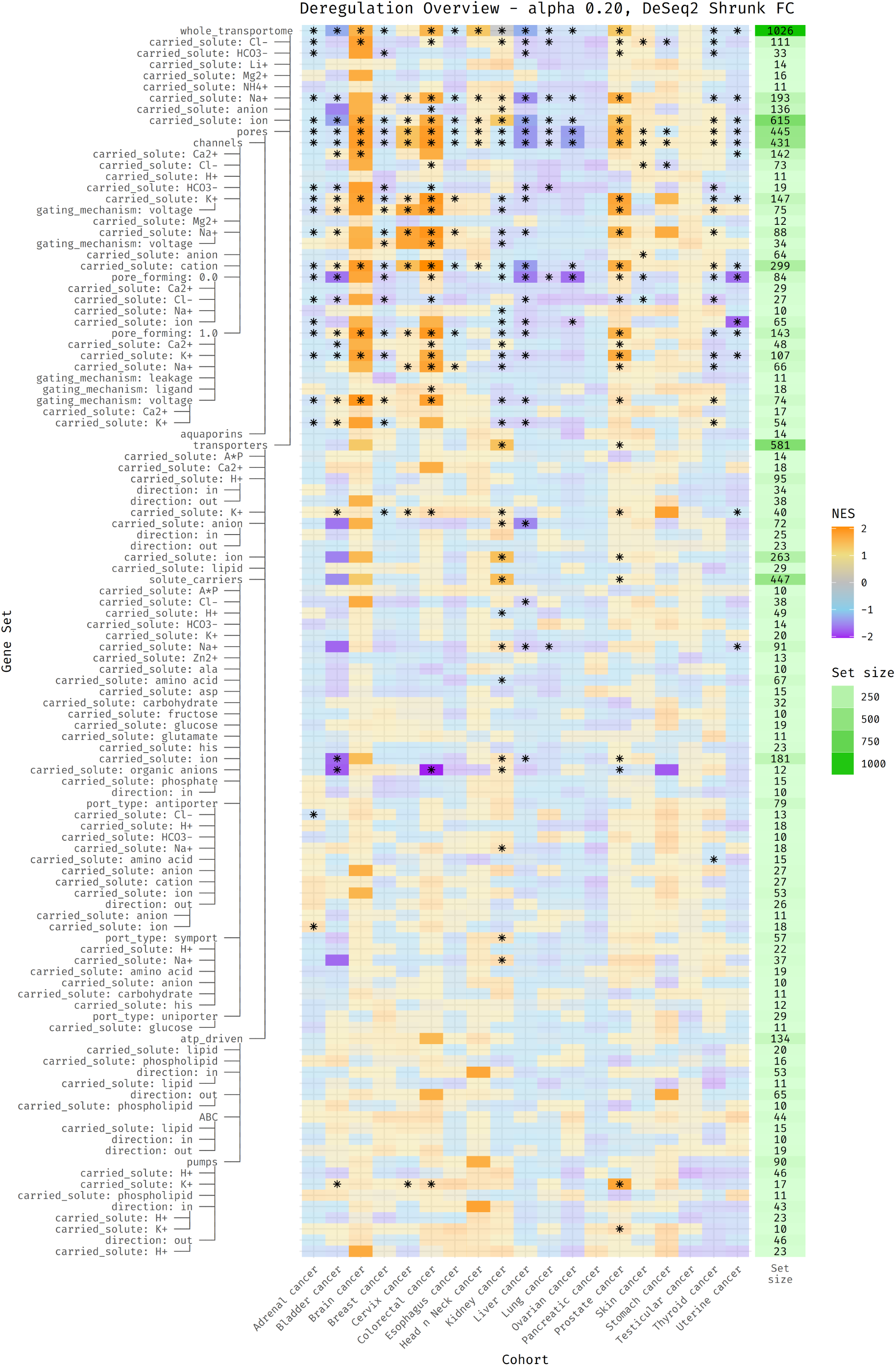
Supplementary deregulation heatmap with metric “deseq_shrinkage”.

**Supplementary Figure J:**
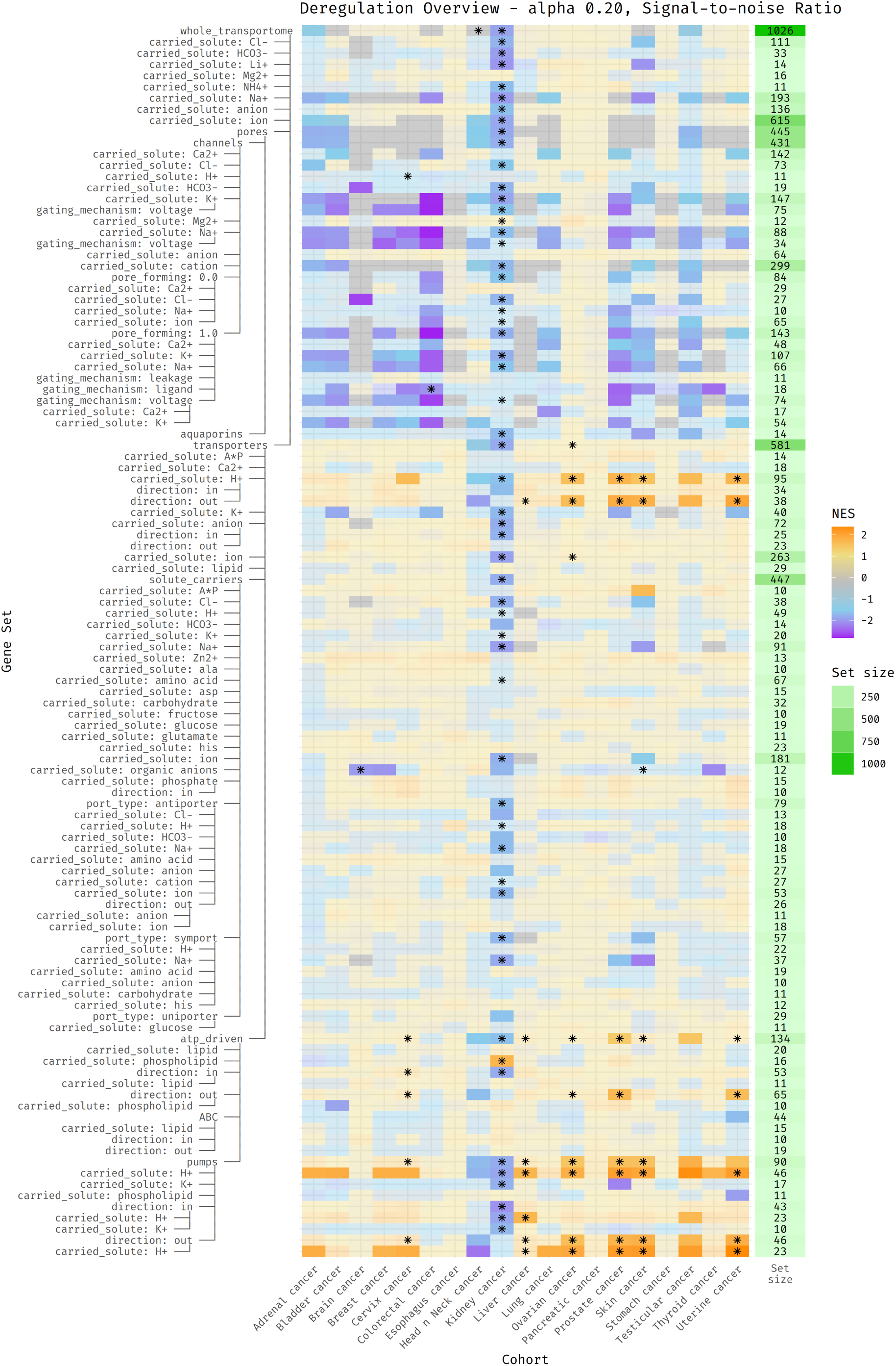
Supplementary deregulation heatmap with metric “s2n_ratio”.

**Supplementary Figure K:**
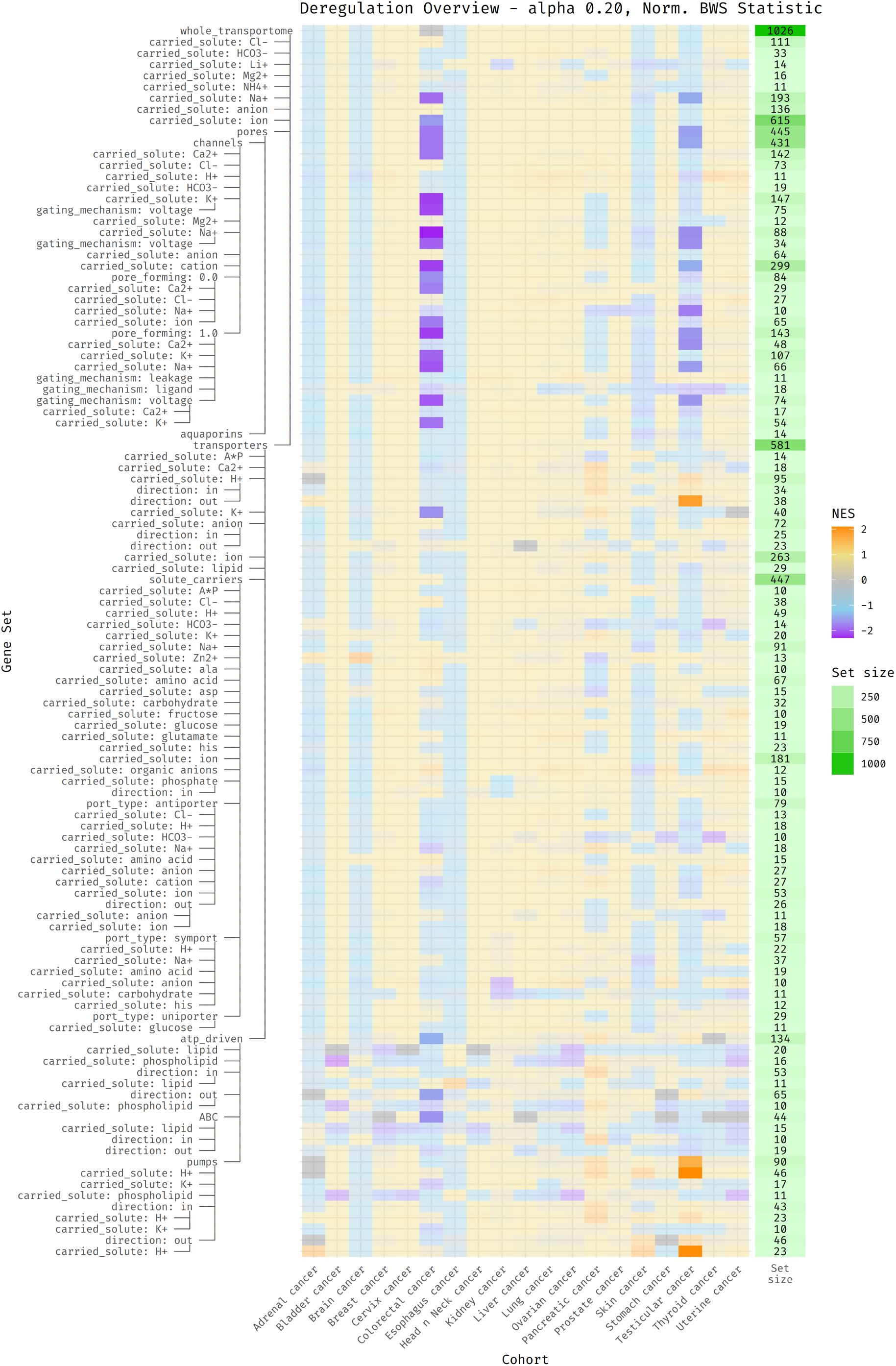
Supplementary deregulation heatmap with metric “norm_bws_test”.

**Supplementary Figure L:**
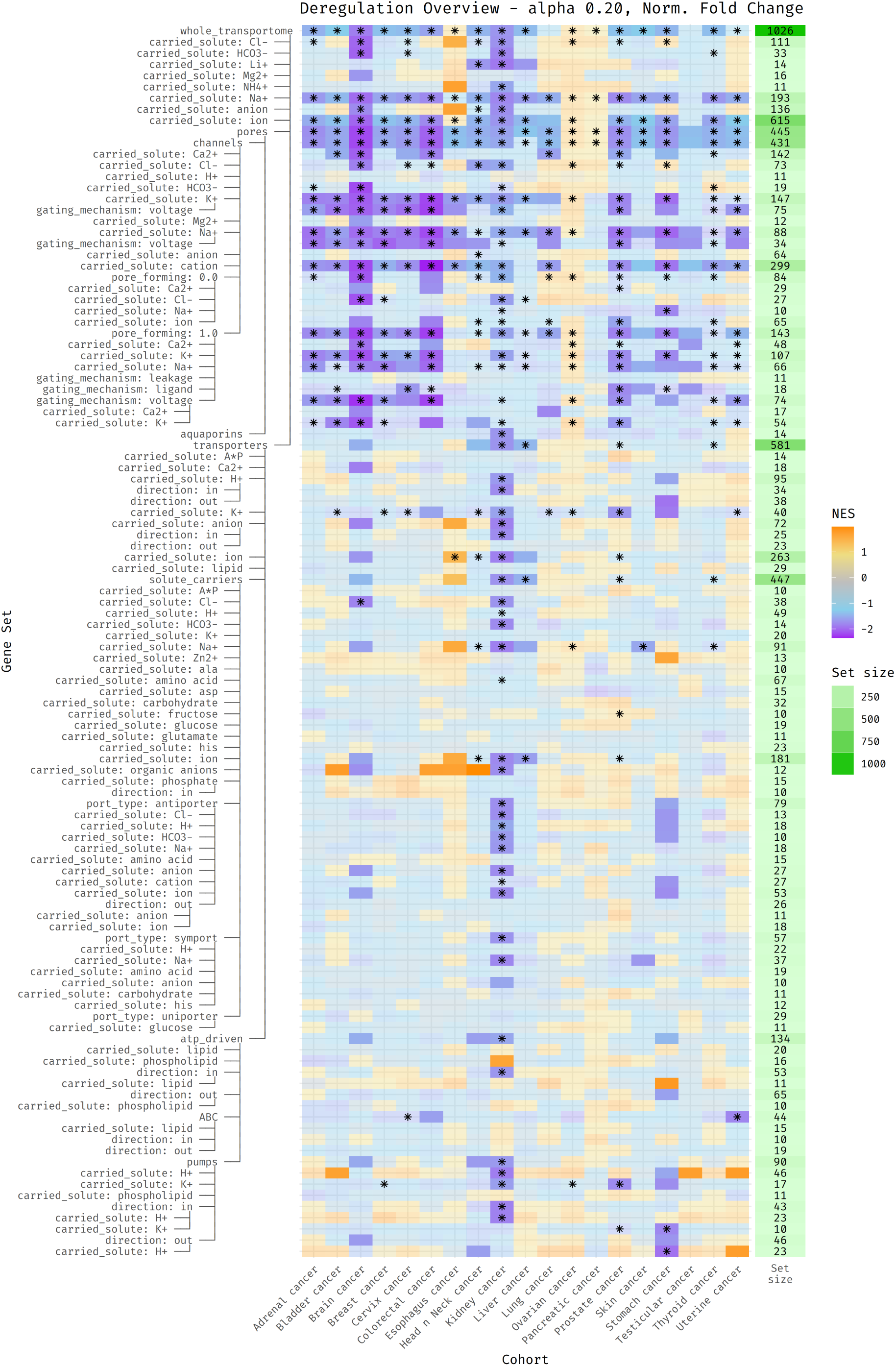
Supplementary deregulation heatmap with metric “norm_fold_change”.

**Supplementary Figure M:**
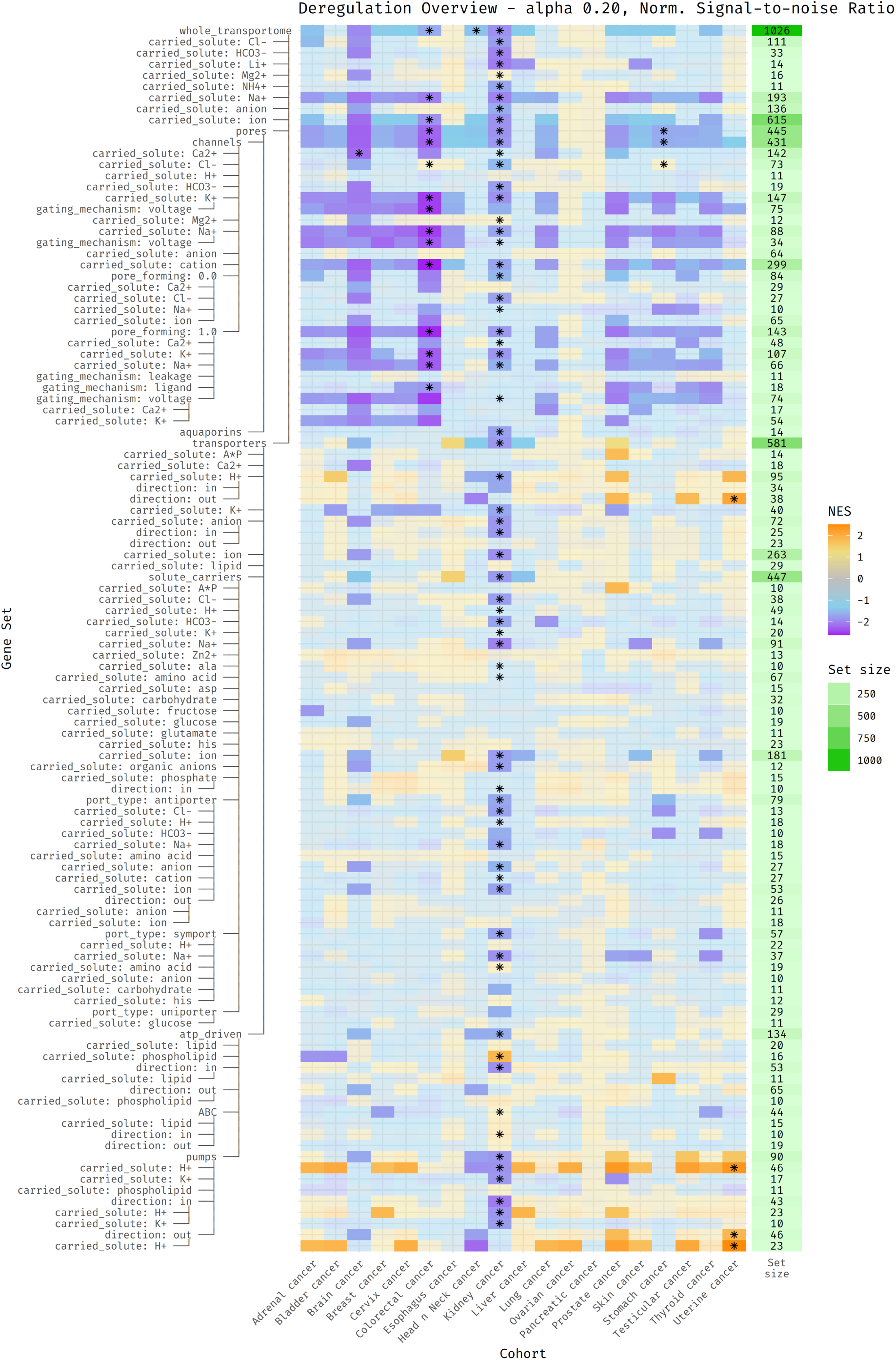
Supplementary deregulation heatmap with metric “norm_s2n_ratio”.

**Supplementary Figure N:**
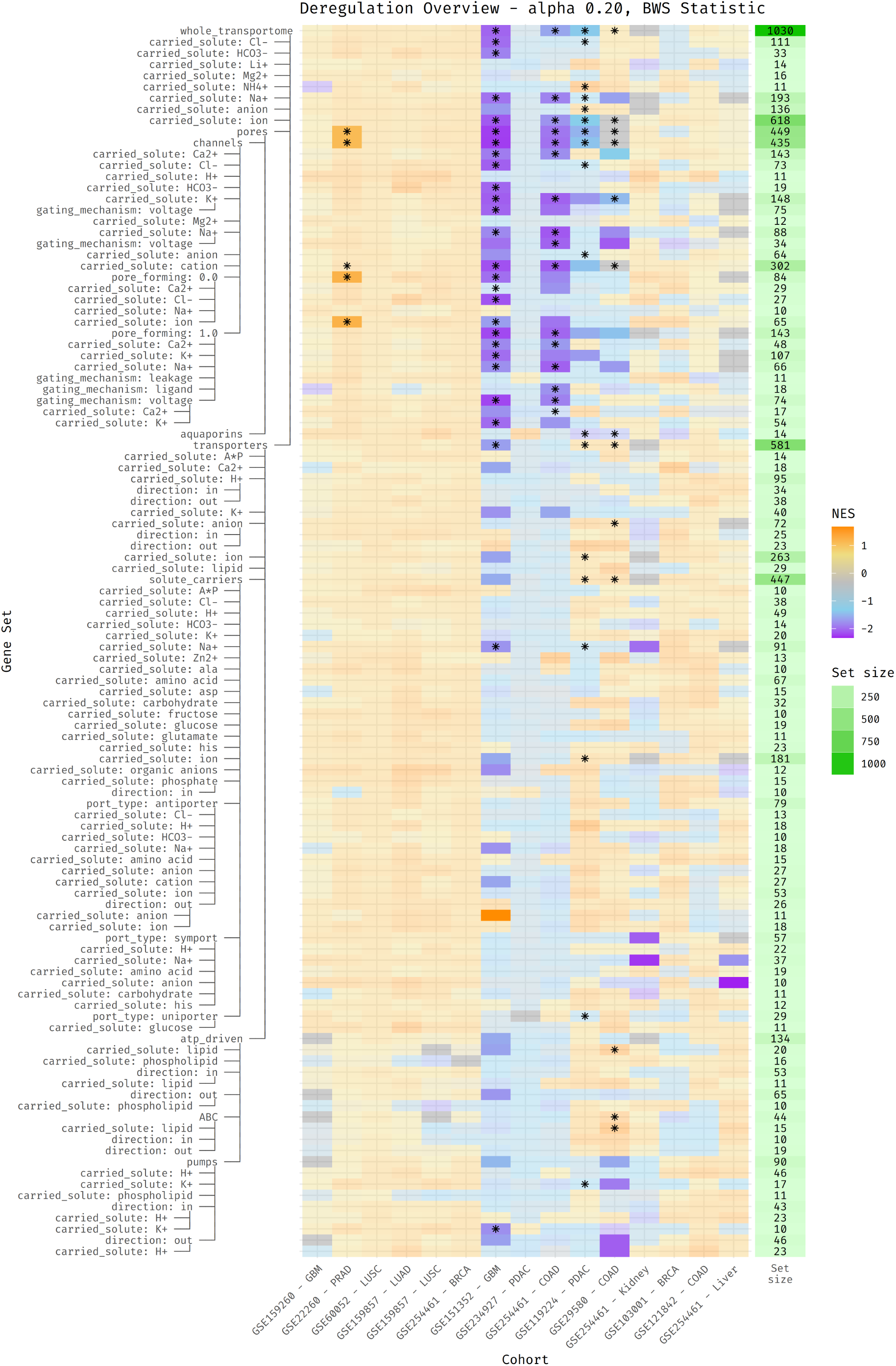
Supplementary deregulation heatmap with metric “bws_test”.

**Supplementary Figure O:**
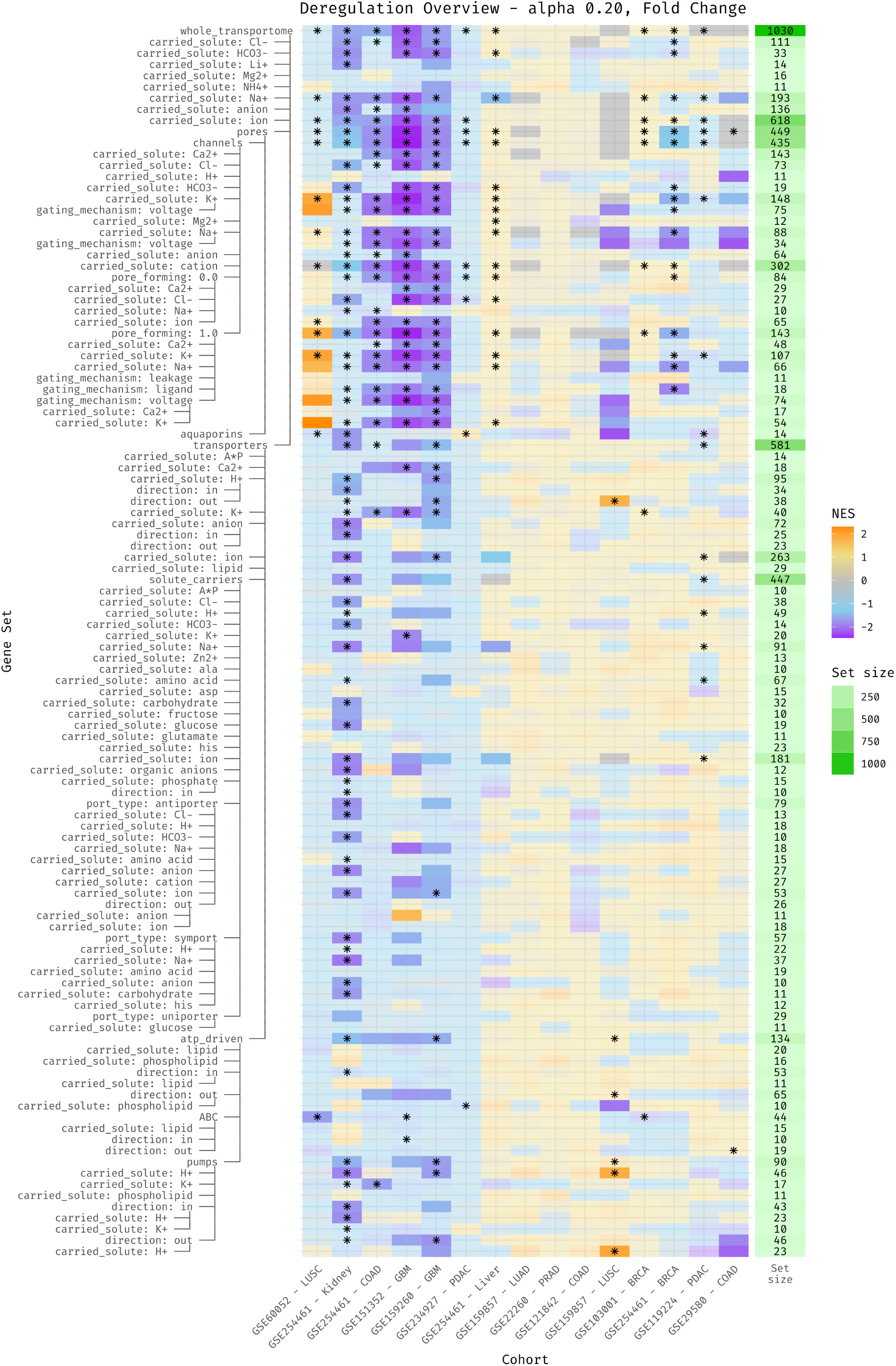
Supplementary deregulation heatmap with metric “fold_change”.

**Supplementary Figure P:**
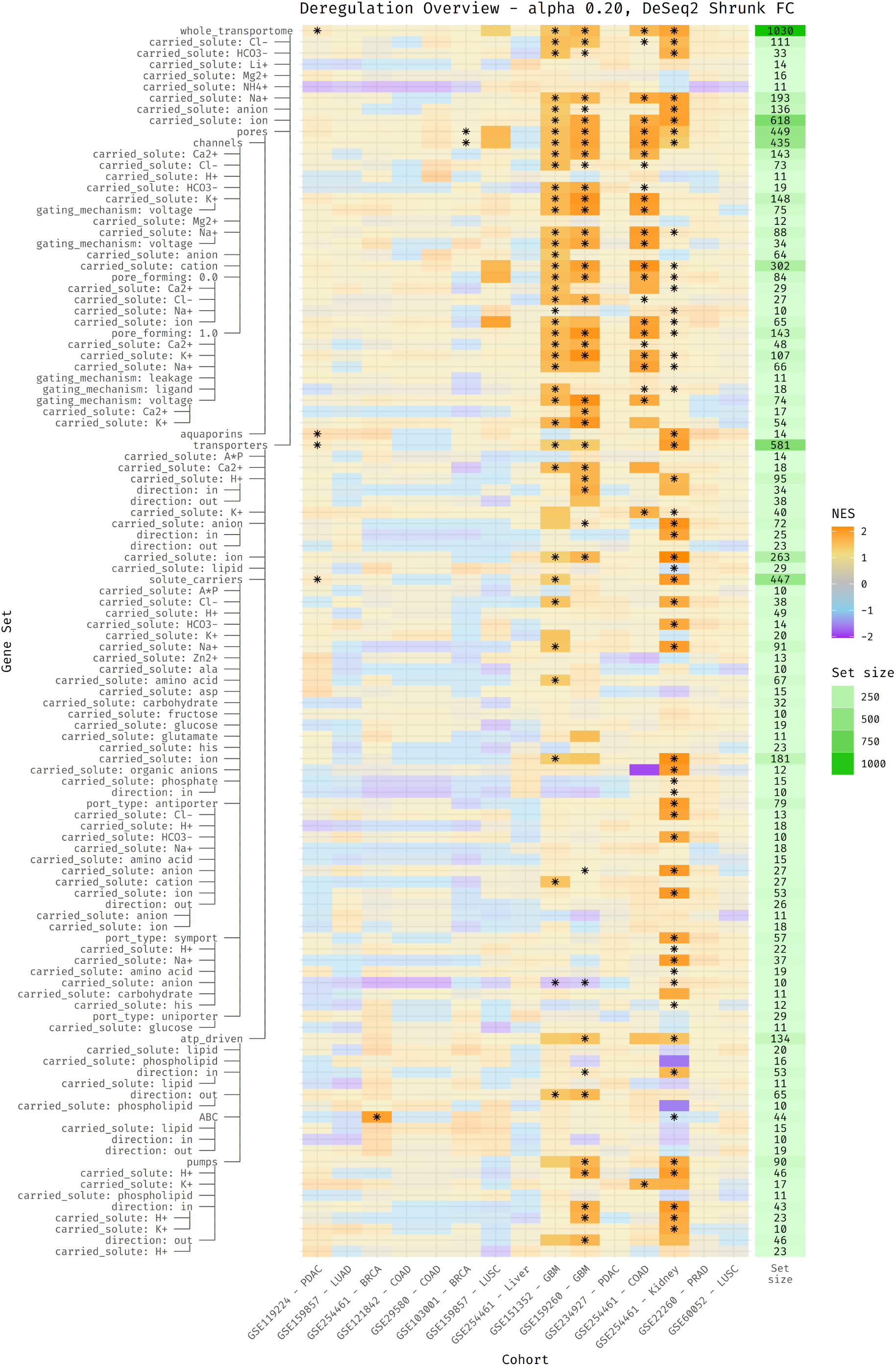
Supplementary deregulation heatmap with metric “deseq_shrinkage”.

**Supplementary Figure Q:**
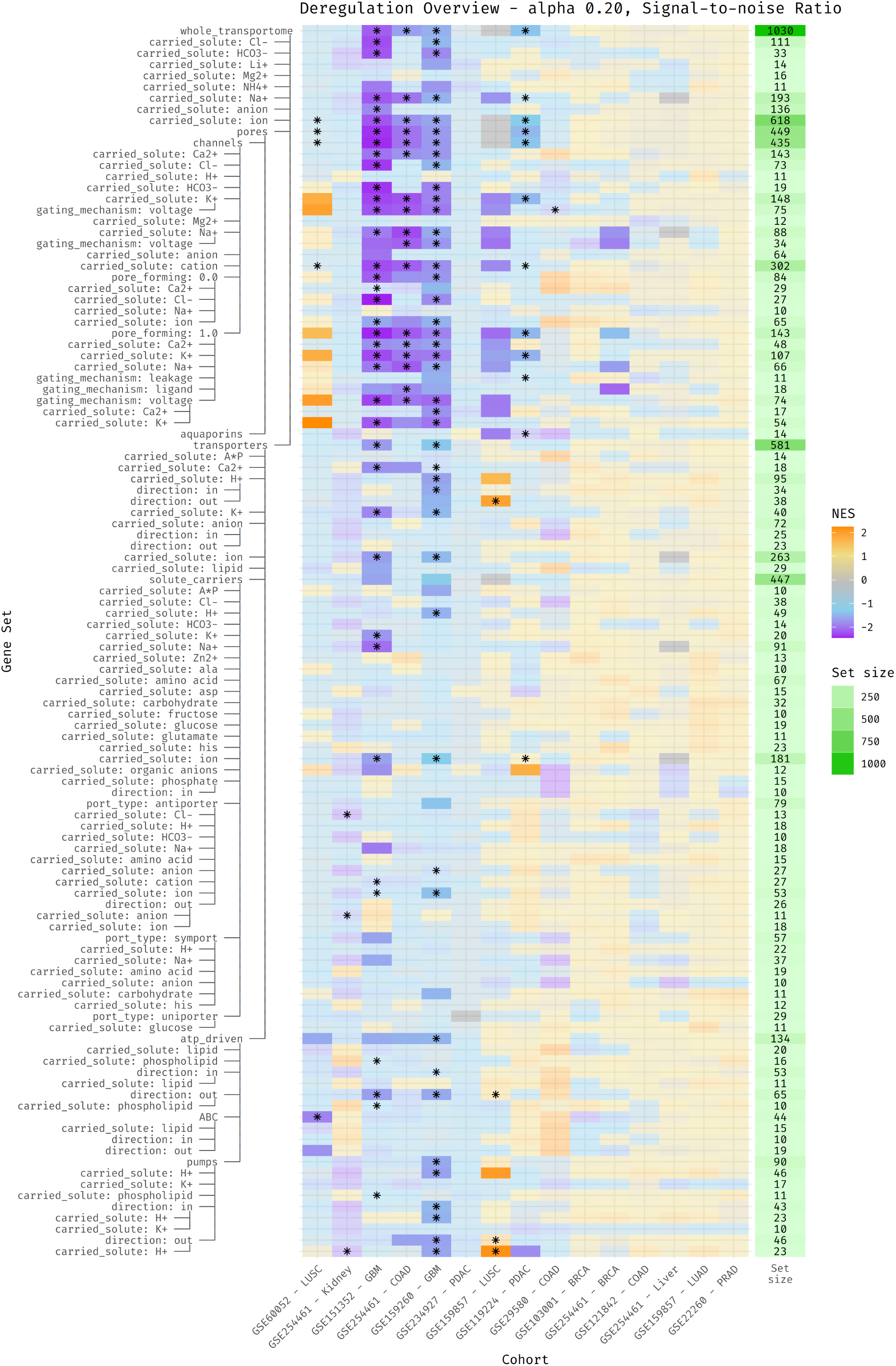
Supplementary deregulation heatmap with metric “s2n_ratio”.

**Supplementary Figure R:**
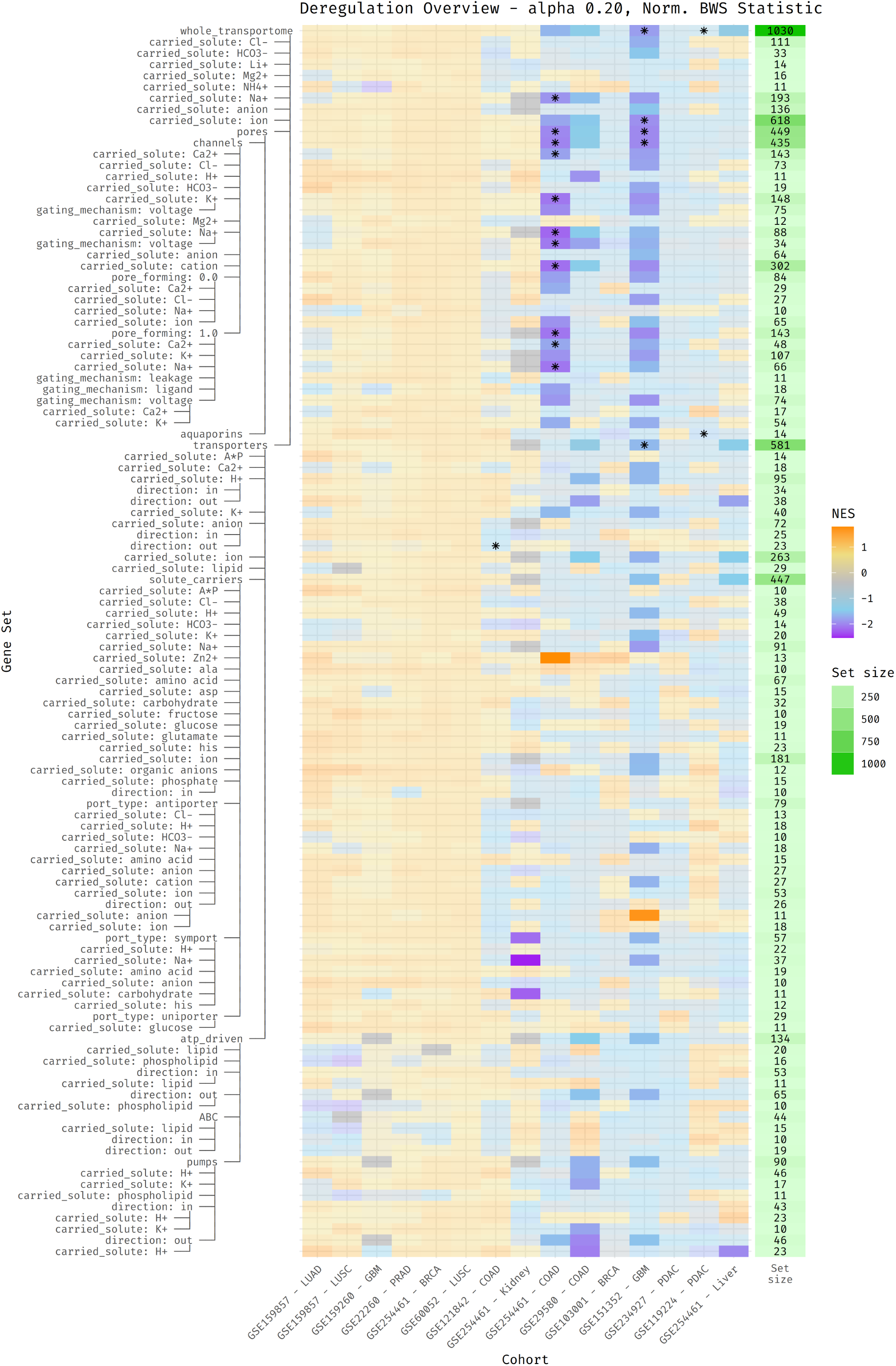
Supplementary deregulation heatmap with metric “norm_bws_test”.

**Supplementary Figure S:**
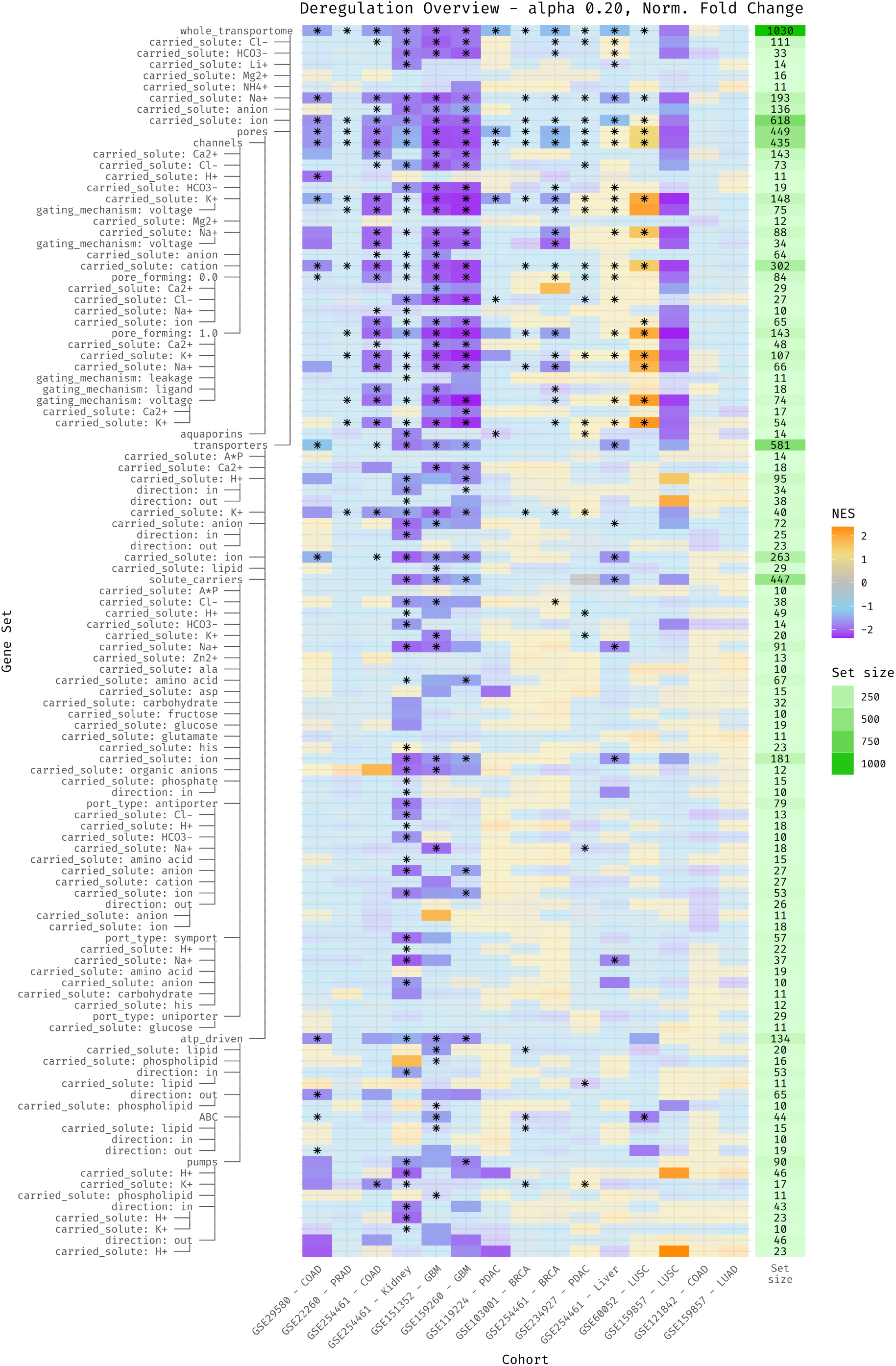
Supplementary deregulation heatmap with metric “norm_fold_change”.

**Supplementary Figure T:**
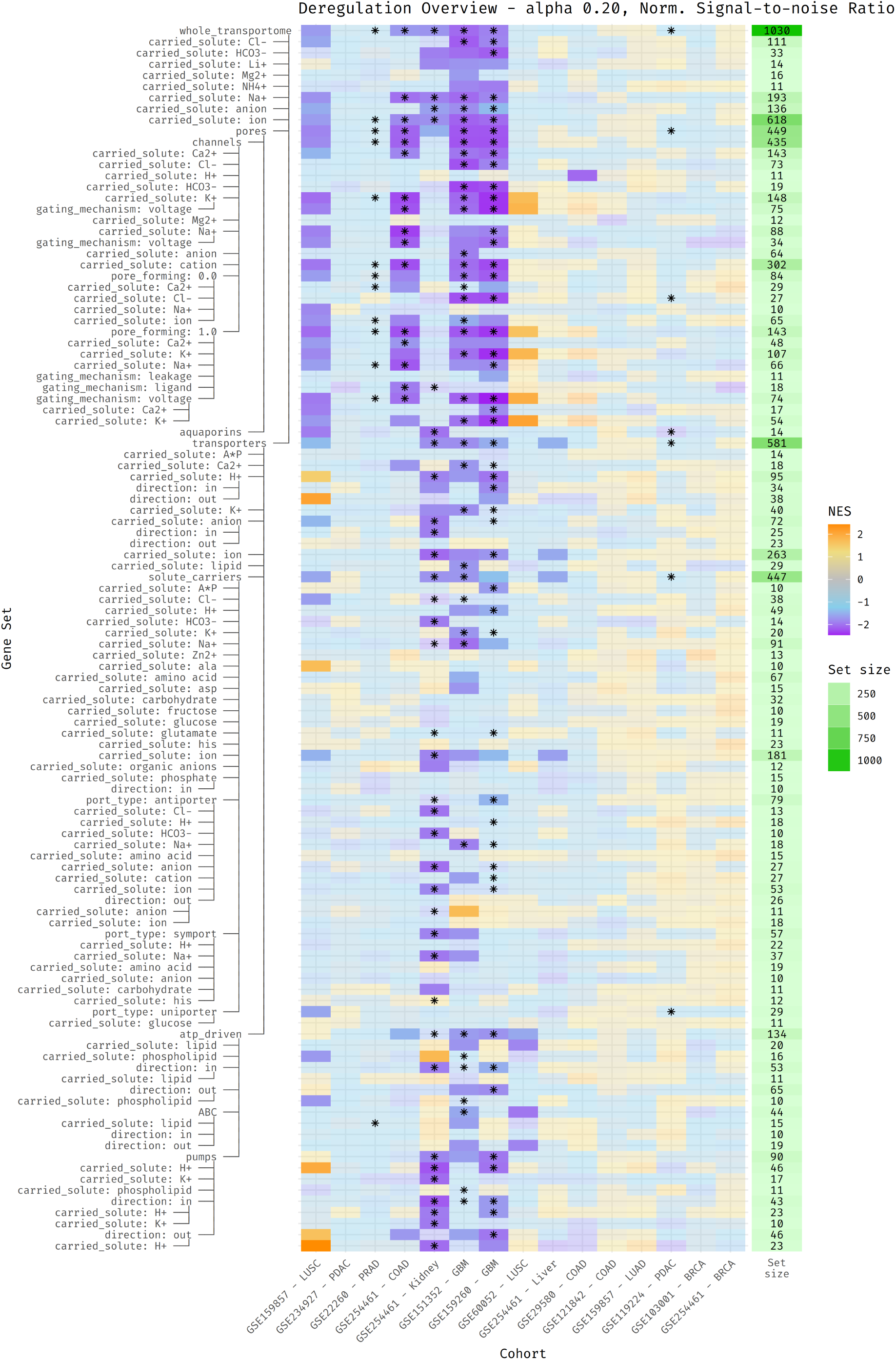
Supplementary deregulation heatmap with metric “norm_s2n_ratio”.

## S.2 Comparison of ranking metrics

To check the congruency of different metrics as computed by `gene_ranker`, we implemented an R package called `compare_ranks` (available on GitHub at TCP-Lab/compare_ranks) which can be used to compare differently ordered lists of items. To do this over two lists, it runs a rolling window of size `n` over both lists. At each step, it checks the number of items in the window of the first list that are also in the same window in the second list. A counter keeps track of the total number of these “hits”. At the end of the run, this total is normalized by the total length of all the windows taken in consideration, resulting in a value between 1 and 0, where 1 is perfect congruency (all items are in the same order as the second list) and 0 (there are no items in the same order).

This score represents the congruency of two lists, but it is dependent on the granularity of the window size: smaller windows check for more granular similarity, while larger windows represent less specific similarities. To avoid having to select a proper window length, the algorithm is run again and again with all possible window sizes, from 1 to the length of the gene set size. This process is very computationally expensive, so we randomly sampled the same subset of 250 genes from each ranking to be compared. This provides computational speed at little cost to the expressiveness of the result.

**Supplementary Figure U:**
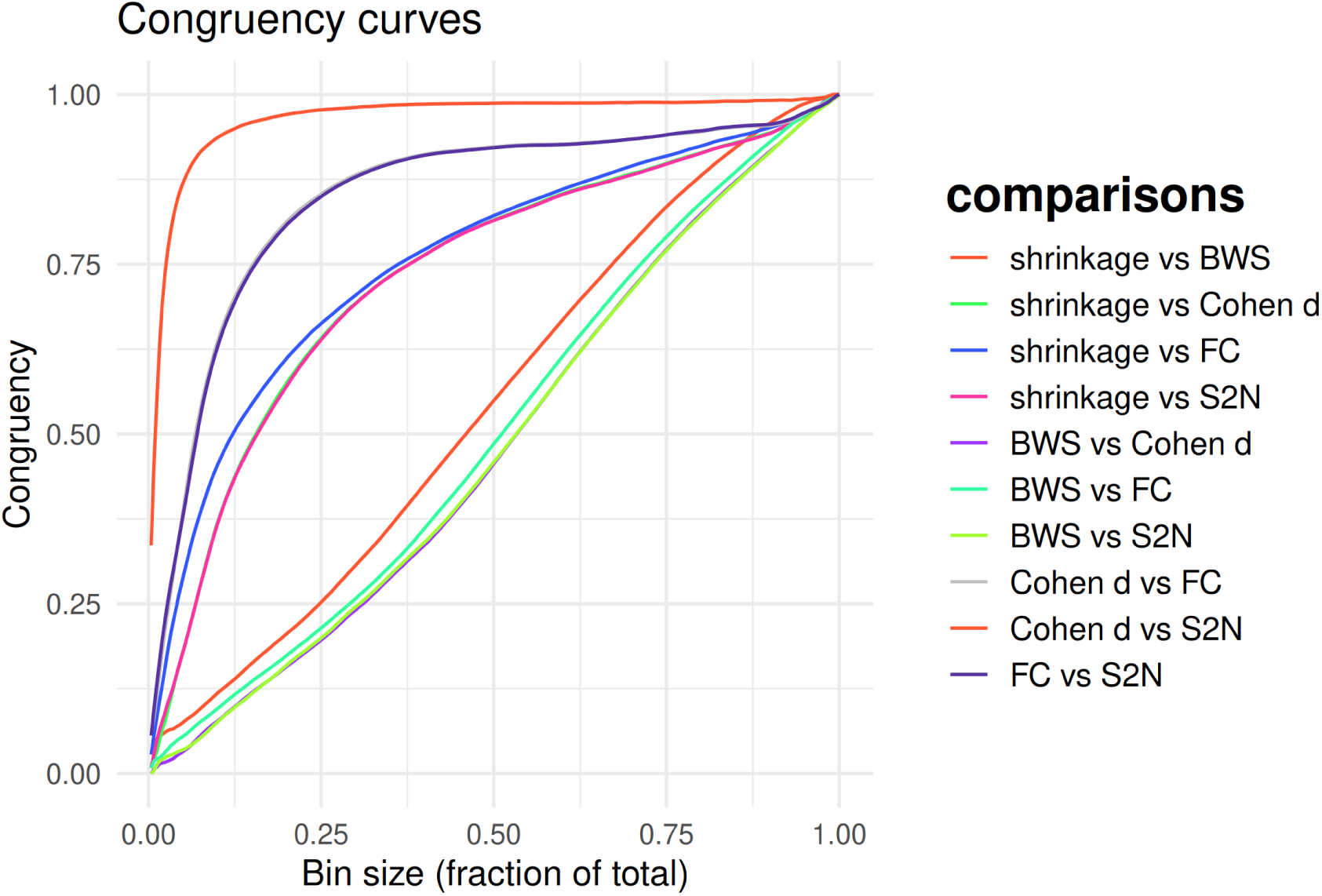
Congruency curves for different metric types. Congruency is a measure between 1 (perfect congruency) and 0 (no congruency). The x axis represents the size of the rolling window used to compute the congruency metric. The area under the curve is proportional to the overall similarity of the two ranked lists. Shrinkage: DeSeq2 shrunk log fold change; BWS: Baumgartner-Weiss-Schindler test statistic; FC: Fold Change; S2N: Signal to noise ratio; Cohen d: Cohen’s D metric. All metrics are normalized as described in the Materials and Methods section.

The congruency of different analyzed metrics can be seen in supplementary figure u. While there was consensus between the simple fold-change and Cohen’s D metrics, the differences were more pronounced with other metrics, such as pydeseq2 shrunk fold change and Baumgartner-Weiss-Schindler test statistic. Relevant to note, the BWS metrics was suggested as one of the best metrics to detect differentially expressed groups with GSEA by Zyla, J., Marczyk, M., Weiner, J. & Polanska, J. [99].

## S.3 Alternate Shared Dysregulation Maps

Figure 4 shows the top 100 Ion Channels and Transporters (ICTs) most-dysregulated genes in all 19 cancer types that were considered.

Here we show identical maps, but including only channels (supplementary figure v) or transporters (supplementary figure w).

**Supplementary Figure V:**
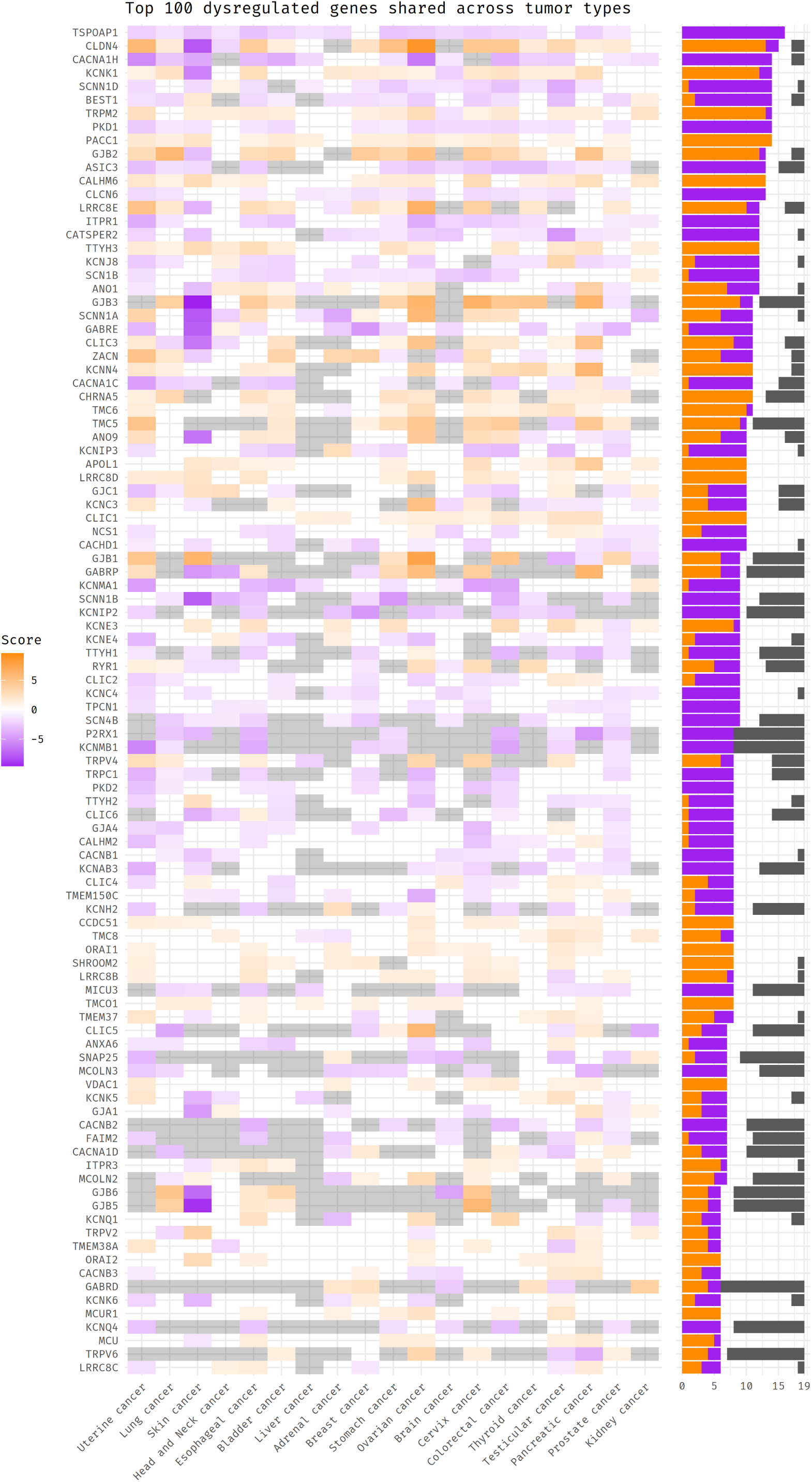
Supplementary “top dysregulation” heatmap considering only channels. All other aspects are identical to Figure 4.

**Supplementary Figure W:**
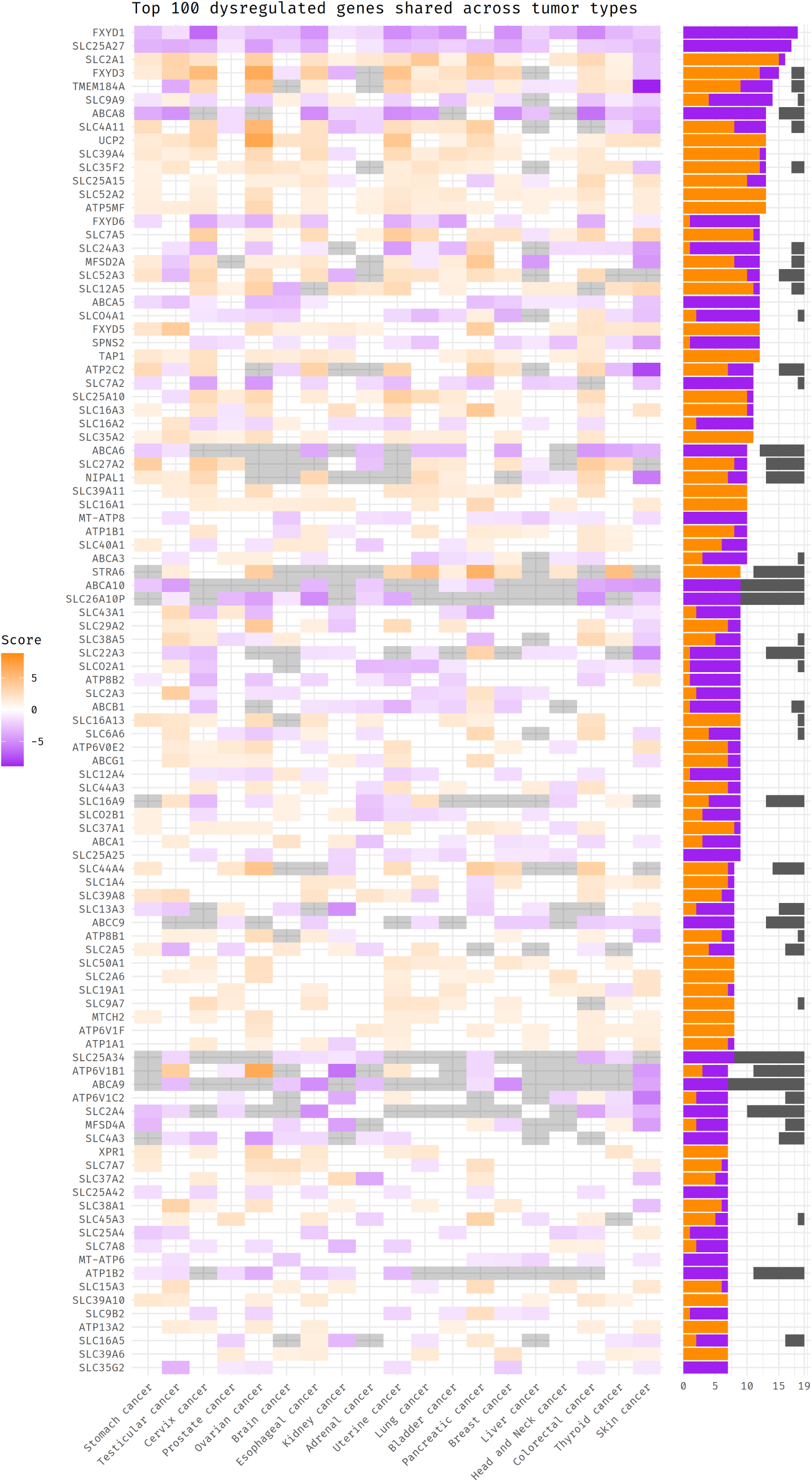
Supplementary “top dysregulation” heatmap considering only trans-porters. All other aspects are identical to Figure 4.

## S.4 Correlation between genesets

To visualize correlation between genesets we calculated the Sørensen–Dice coefficient between every pair of genesets that survived pruning. These coefficients are plotted in supplementary figure x.

**Supplementary Figure X:**
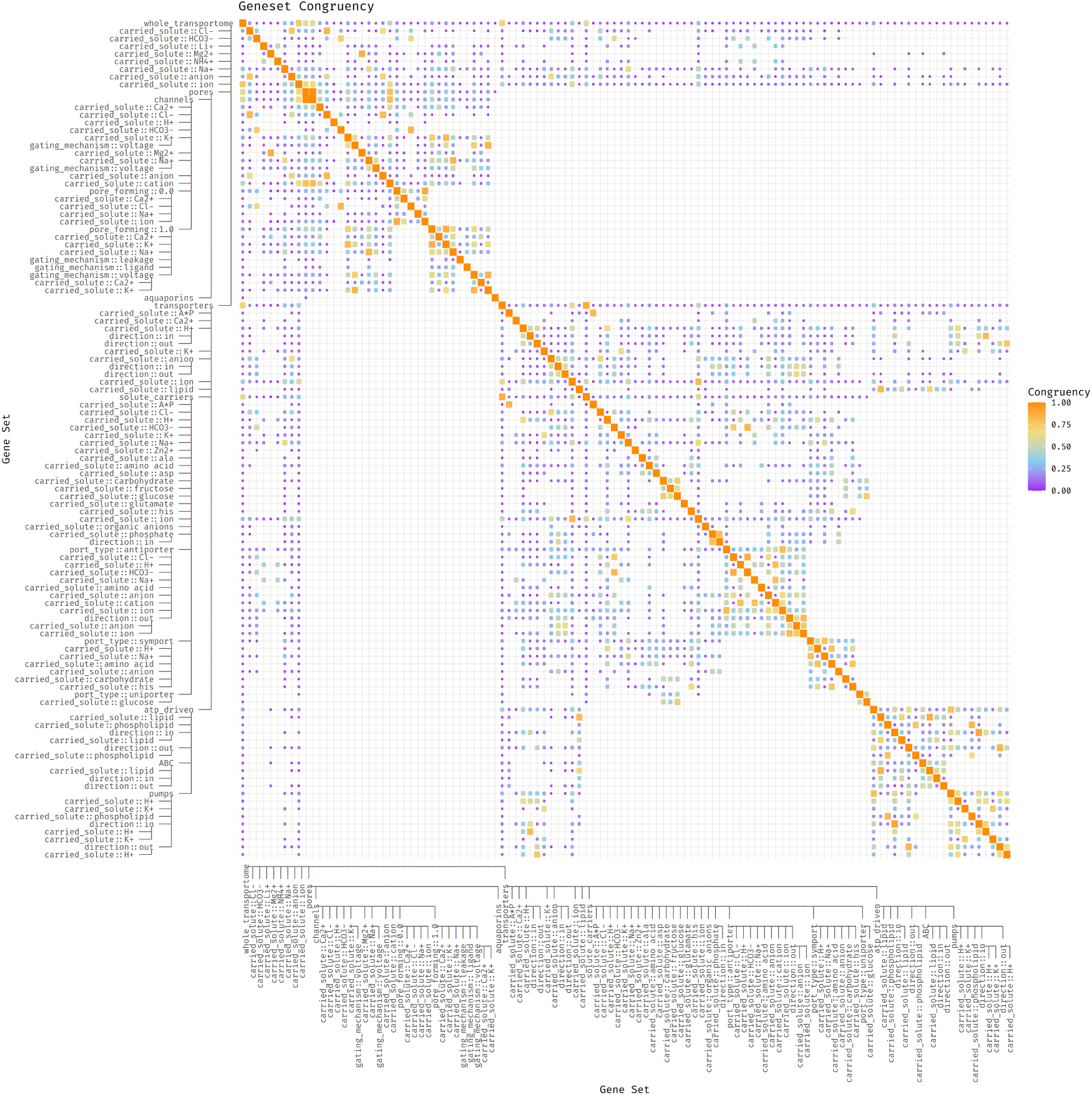
Heatmap of genesets congruency. Displayed is the Sørensen–Dice coefficient described in Section 5 computer for every pair of genesets after pruning. Color and size of the heatmap square are proportional to the value of the coefficient. A coefficient of 1 means identical genesets. A congruency of 0 means completely different genesets. Overall correlation between parent and child notes is kept low thanks to the pruning approach detailed in the Materials and Methods section.

